# Type 1 and type 2 dendritic cell subsets cooperate to maintain intestinal immune tolerance via integrin αvβ8-mediated TGF-β activation

**DOI:** 10.64898/2026.02.26.707908

**Authors:** Sébastien This, Delphine Brichart-Vernos, Venla Anniina Väänänen, Carine Rey, Véronique Barateau, Tanguy Fenouil, Urs Michael Mörbe, Lucie Daniel, Stephen L. Nishimura, Stéphanie Graff-Dubois, Olivier Thaunat, William W. Agace, Helena Paidassi

## Abstract

The gastrointestinal tract is a unique immunological environment where the host must balance tolerance to commensal microbes with defense against pathogens. A critical mechanism for maintaining this balance is the peripheral conversion of naïve T cells into regulatory T cells (pTregs), a process that depends on the TGF-β cytokine, which is produced in a latent form and must be activated. While the activation of latent TGF-β relies on the membrane-bound αvβ8 integrin, the precise cellular subset(s) responsible for this essential process have yet to be clearly defined. Conventional dendritic cells (cDCs), which migrate from the intestinal lamina propria to the gut-draining mesenteric lymph nodes (MLN), have long been considered the primary antigen-presenting cells (APCs) responsible for αvβ8-mediated TGF-β activation and pTreg induction. However, recent studies have challenged this paradigm by highlighting a new family of rare RORγt-expressing APCs, able to induce pTreg via αv integrins, raising questions about the *in vivo* role of cDCs in the maintenance of mucosal immune homeostasis.

Using a β8 integrin gene reporter mouse model (*Itgb8-IRES-tdTomato*) combined with single-cell profiling, we comprehensively mapped *Itgb8*-expressing APCs in the MLN. We show that cDCs, in particular migratory type 1 (cDC1) and type 2 (cDC2), constitute the predominant *Itgb8^TdTomato^*^+^ cells, both in neonatal and adult mice. Through cDC subset-specific β8 knockout models, we demonstrate that both cDC1 and cDC2 are required for optimal pTreg generation. Loss of β8 integrin in either subset led to a partial reduction in pTreg, while combined deletion resulted in profound pTreg loss and spontaneous colitis. Importantly, these effects were independent of RORγt⁺ APC populations, including ILC3s and Thetis cells.

These findings resolve longstanding questions about the identity of key APCs driving pTreg induction in the MLNs. They demonstrate that cDC1 and cDC2 are non-redundant, essential mediators of pTreg induction and intestinal immune tolerance. Although different populations of RORγt⁺ APCs may contribute in specific contexts, such as early development, infection, or in the prevention of allergic disease, cDCs remain one of the primary guardians of intestinal immune homeostasis in response to microbiota.

## INTRODUCTION

The gastrointestinal tract is home to a diverse population of commensal bacteria that provide significant benefit to the host, including nutrient processing and immune education. Yet, it also represents a major entry point for pathogens. The mucosal immune system must balance robust protection against infection with tolerance towards beneficial commensals, maintaining homeostasis. Disruptions of this delicate equilibrium is believed to significantly contribute to the pathogenesis of inflammatory bowel disease (IBD)^1,2^.

Forkhead box P3 (FoxP3)-expressing regulatory T cells (Treg) are key mediators of immune tolerance in the intestine^3^. These cells can be categorized into two major subsets: thymically derived Tregs (tTreg), which develop in the thymus and prevent systemic autoimmunity, and peripherally induced (pTreg), which arise from naïve T cells in the periphery and play a pivotal role in maintaining tolerance to foreign intestinal antigens, including those from commensal microbes. Among pTreg, RAR-related orphan receptor gamma (RORγt) expression further differentiates pTreg according to function. RORγt-expressing Treg provide tolerance to the microbiota while RORγt^-^ Treg are preferentially involved in tolerance to food antigens^4^. While Treg deficiency causes widespread autoimmunity, a selective impairment in pTregs generation is associated with intestinal inflammation and IBD, in both mice and humans^5–8^.

Conventional dendritic cells (cDCs) have long been recognized as the primary antigen-presenting cells (APCs) responsible for pTreg induction in the mesenteric lymph nodes (MLNs). In the lamina propria (LP) of the small and large intestine, cDCs express the αE(CD103)β7 integrin. Upon activation, CD103^+^ cDCs migrate to the MLNs where they present intestinal antigen to naïve T cells and preferentially promote pTreg differentiation, in a Transforming Growth Factor beta (TGF-β)-dependent manner^9–11^. TGF-β is secreted in latent inactive form and must be activated to bind to its receptor and exert its immunomodulatory effects^12^. αvβ8 integrin-mediated TGF-β activation is a key mechanism for the regulation of TGF-β bioavailability in the immune system^12^. Prior studies, including our own, demonstrated that while αv integrin is ubiquitously expressed, the CD103^+^ MLN cDCs preferentially express the β8 integrin gene (*Itgb8*)^13–15^. *In vitro*, αv or β8 integrin deficient CD103^+^ cDCs display impaired TGF-β activation and a reduced capacity to induce Treg^13,14^. Mice with a specific deletion of either αv or β8 in CD11c^+^ cells develop spontaneous colitis and show defects in pTreg generation, emphasizing the importance of αvβ8-mediated TGF-β activation by cDCs in intestinal homeostasis^16,17^.

For many years, this model formed the cornerstone of our understanding of peripheral tolerance: that MLN CD103⁺ cDCs were the dominant APCs driving pTreg differentiation. CD103^+^ cDCs are themselves heterogeneous and comprise two main subsets, type 1 (cDC1) and type 2 (cDC2), distinguished by surface markers and transcriptional factor dependencies, and which have different functions (reviewed in ^18^). cDC1 (CD11b^-^) dependent on IRF8 and BATF3 are specialized in promoting CD8^+^ cytotoxic and CD4^+^ T-helper 1 (Th1) responses to virus and intra-cellular bacteria^19,20^. In contrast, cDC2 (CD11b^+^) dependent on IRF4, preferentially drive Th2, Th17 and T follicular helper (Tfh) responses to extracellular pathogens^21–26^. With respect to pTreg induction, recent transcriptomic and functional analyses had suggested MLN cDC1 may be particularly proficient at promoting pTreg differentiation through αvβ8-mediated TGF-β activation^27,28^. These findings reinforced the long-standing paradigm that cDCs are central to the establishment of intestinal tolerance. However, despite compelling *in vitro* data, the *in vivo* relevance of these findings remained unresolved.

This framework was fundamentally challenged by a series of studies published in 2022, which highlighted a novel family of RORγt-expressing APCs, as potential inducers of intestinal pTreg^29–31^. These RORγt^+^ APCs include group 3 innate lymphoid cells (ILC3s), extrathymic AIRE-expressing cells (eTACs), Janus cells (JCs) and Thetis cells^29–31^. Like cDCs, these RORγt^+^ APCs also express αvβ8 integrin and have been implicated in the maintenance of intestinal immune tolerance. Strikingly, these findings not only introduced new APC subsets involved in pTreg generation but also proposed that cDCs may not play a significant role in this process at all, suggesting a dramatic paradigm shift away from the long-held dogma that positioned cDCs at the centre of mucosal tolerance. However, it is critical to note that, in the studies which concluded to the lack of necessity of cDC subsets in the maintenance of intestinal immune homeostasis, *ClecSa^Cre^* mice were used to delete MHC-II specifically in cDCs^29,31^. Yet, while *ClecSa^Cre^* targeting is indeed highly specific to cDCs, it is now well established that this system suffers from limited recombination efficiency, leaving 70-80% of cDCs functionally intact^32,33^. As such, conclusions drawn from these models may underestimate the true contribution of cDCs in pTreg induction.

In this study, we re-examined the role of αvβ8 integrin-expressing APCs in murine MLNs, with a focus on the specific contribution of cDC1 and cDC2. We found that migratory cDC1 and cDC2 are the predominant *Itgb8*-expressing APCs in the MLNs of both neonatal and adult mice. Targeted deletion of β8 integrin in either cDC1 or cDC2, without impacting RORγt^+^ APCs, resulted in a significant reduction in intestinal pTreg generation. Strikingly, only when β8 integrin was deleted in both RORγt^-^cDC1 and cDC2 subsets did we observe spontaneous colitis and a profound loss of intestinal pTreg. These findings demonstrate that both cDC1 and cDC2 subsets play complementary and essential roles in αvβ8-mediated activation of TGF-β, thereby driving pTreg differentiation and maintaining intestinal immune homeostasis. Finally, we show that human migratory DCs also express β8 integrin-encoding gene (*ITGB8*), highlighting the potential translation relevance of our findings.

Together, our data clarify a central question in mucosal immunology regarding the identity and relative importance of APC subsets in pTreg induction. While RORγt⁺ APCs, particularly Thetis cells, may contribute during a narrow developmental window, migratory cDC1 and cDC2 remain essential for maintaining peripheral tolerance through adulthood, owing to their unique capacity for TGF-β activation via αvβ8 integrin. Our findings underscore the functional cooperation among APC subsets and highlight the evolutionary importance of redundant yet complementary tolerance mechanisms to safeguard intestinal homeostasis and prevent inflammatory disease.

## RESULTS

### Migratory cDCs are the most abundant MHCII^+^ APCs expressing the TGF-β-activating integrin αvβ8

While the relative contributions of cDCs and RORγt^+^ APCs in maintaining intestinal immune tolerance remain unclear, previous studies suggest that both cell types require αvβ8 integrin to induce pTreg via TGF-β activation. Integrins are obligate heterodimeric trans-membrane receptors, composed of one α and one β subunits. Although αv integrin is broadly expressed, TGF-β activation, and thus pTreg differentiation, is regulated specifically at the level of β8 integrin gene (*Itgb8*) expression in MLN CD11c^+^ APCs^13,14,28^.

To uncover the full spectrum of APCs competent for αvβ8-mediated TGF-β activation, we leveraged our recently developed *Itgb8-IRES-tdTomato* (*Itgb8^tdTomato^*) reporter mouse model, which enable single-cell resolution analysis of *Itgb8* gene expression^34^. We enriched APC populations by depleting monocytes, granulocytes, T, B and NK cells (Lineage^+^ cells) from pooled MLNs of two-week-old neonate and sixteen-week-old adult *Itgb8^tdTomato^*mice. We then sorted live CD45^+^Lin^-^MHCII^+^ Itgb8(Tomato)^+^ cells (hereafter, ‘Itgb8(Tomato)^+^ APC’) for single-cell RNA sequencing (scRNA-seq). As a reference, we also sorted total CD45^+^Lin^-^MHCII^+^ APCs (‘total APCs’) from age-matched control mice to capture the broader heterogeneity of MLN APCs (**Figure 1A** and **Supplemental Figure S1**).

**Figure 1.**
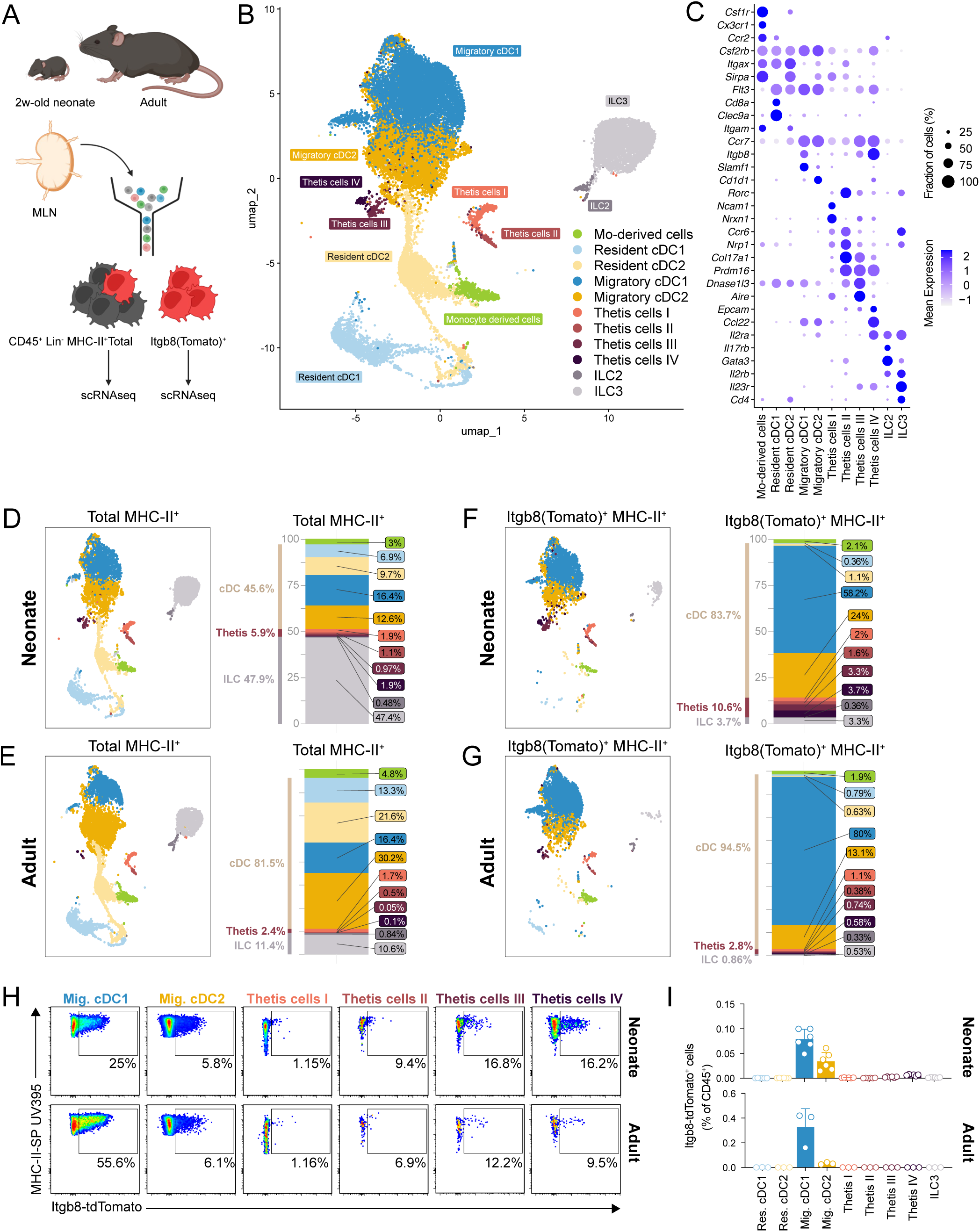
Migratory cDC1 and cDC2 are the predominant αvβ8-expressing APCs in the MLN of neonatal and adult mice. **A.** Schematic of scRNA-seq of total and Itgb8(Tomato)+ Lin–MHCII+ cells from the MLNs of 2-week-old neonatal or adult *Itgb8-IRES-tdTomato* (*Itgb8^tdTomato^*) reporter mice. **B**. Uniform manifold approximation and projection (UMAP) visualization of 24,403 cells obtained from scRNA-seq of MLN APCs, combining data from sorted total MHC-II^+^ and Itgb8(Tomato)^+^ MHC-II^+^ cells from 2-week-old and adult mice for joint clustering, colored by cluster annotation. **C**. Dot plot showing the expression of canonical DC, Thetis and ILC marker genes on total MLN APC subsets. **D-E** UMAP plot and stacked bar plots comparing the proportion of each cluster in ‘total’ APCs from 2-week-old neonatal mice (8,278 cells) (**D**) and adult mice (9,365 cells) (**E**). **F-G** UMAP plot and stacked bar plots comparing the proportion of each cluster in ‘Itgb8(Tomato)^+^ APCs from 2-week-old neonatal (2,815 cells) (**F**) and adult mice (3,945 cells) (**G**). **H-I**. Itgb8(Tomato) expression in each APC subset (gated as in Supplemental Figure 4) from the MLNs of 2-week-old neonatal and adult (15 to 19-week-old) *Itgb8^tdTomato^* reporter mice. (**H**) Representative flow cytometry plots show tdTomato expression in neonatal (top; n = 6) and adult (bottom; n = 3) samples. Data are shown as concatenated events from pooled biological replicates. (**I**) Proportion of Itgb8(Tomato)^+^ cells for the indicated population in total CD45^+^ cells. Error bars represent S.D.

In total, we profiled eight scRNA-seq samples from neonatal and adult mice across four genetic models. All datasets were integrated and clustered together to generate a unified transcriptional atlas of MLN APCs. For clarity, our initial analysis focuses on total and Itgb8(Tomato)⁺ APCs from *Itgb8^tdTomato^*and control mice, which served as the foundation for defining APC subsets. The four additional samples—derived from Cre-reporter fate-mapping models—are introduced in subsequent sections. Following quality control filtering, we retained transcriptional profiles for 38,534 high-quality cells. Unsupervised clustering revealed eight transcriptionally distinct clusters, visualized by Uniform Manifold Approximation and Projection (UMAP) (**Figure 1B-C**, **Supplemental Figure S2A**). Two minor contaminant clusters were excluded: cluster_6 (B cells) and cluster_7 (glial cells).

The remaining six clusters were stratified based on *Rorc* expression. Four *Rorc*-negative clusters (cluster_0, cluster_1, cluster_2, and cluster_4) comprised classical APC subsets. Cluster_4 was further resolved into two subclusters: cluster_4_s0 and cluster_4_s1. Based on canonical markers, cluster_4_s1 (*Cx3cr1*, *Ccr2*) was annotated as monocyte-derived cells (**Supplemental Figure S2B**). The remaining four *Rorc*-negative clusters represented conventional DCs (cDCs; *Flt3 Csf2rb*), which further separated into two major subtypes: lymph node ‘resident’ (Cluster_2 and cluster_4_s0; *Itgax*^hi^ *Ccr7*^Low^) and ‘migratory’ intestinal cDCs (Cluster_0 and cluster_1; *Itgax*^Low^ *Ccr7*^Hi^). These were further classified into cDC1 (*Cd8α^+^ClecSa^+^* for resident cDC1 and *Slmaf1^+^* for migratory cDC1) and cDC2 (*Itgam^+^Sirpa^+^*for resident cDC2 and *Cd1d1^+^* for migratory cDC2) (**Figure 1B-C**, **Supplemental Figure S2A-B**).

The two remaining clusters (cluster_3 and cluster_5) expressed *Rorc*, indicative of RORγt^+^ APCs. This is an emerging field with overlapping nomenclature across studies—including Janus cells (JCs), extra-thymic Aire-expressing Cells (eTACs), Thetis cells, PRDM16^+^ tolerizing DCs (tolDCs) and RORγt^+^ DCs ^29–31,35–39^. We adopt the Thetis nomenclature here, while acknowledging its ongoing evolution. To further characterize these clusters, we increased the clustering resolution, which revealed two distinct subclusters within cluster_3, and four within cluster_5 (**Supplemental Figure S2B**). We cross-referenced their transcriptomic profiles with published datasets^29,31^ to annotate these six *Rorc*-expressing populations. The cluster_3 subclusters corresponded to ILC2 (cluster_3_s0; *Il17rb* and *Gata3*) and ILC3 (cluster_3_s1; *Il23r* and *Cd4*) (**Supplemental Figure S2C-D**). The cluster_5 subclusters were assigned to the four subsets of Thetis cell. We detected, *Aire*-expressing Thetis I (cluster_5_s0; *Ncam1*^+^ and *Nrxn1*^+^) (also known as eTAC1, JC1 or RORγt^+^ DC I) and Thetis III cells (cluster_5_s1; *Dnase1l3*^+^) (equivalent to eTAC3, JC2 or RORγt^+^ DC III), as well as Thetis II (cluster_5_s2) marked by expression of *Ccr6* and *Nrp1* (equivalent to eTAC2 or RORγt^+^ DC II), and Thetis IV (cluster_5_s3; *Epcam*^+^, *Ccl22*^+^ and *Il2ra*^+^) (equivalent to RORγt^+^ DC IV) predominantly found in neonates (**Figure 1B-C**, **Supplemental Figure S2E-F**).

Now looking at individual samples, the ‘total APC’ dataset enabled comprehensive identification of 11 distinct APC subpopulations within MLN, each characterized by unique transcriptional signatures (**Figure 1B-C**, **Supplemental Figure S3A-B**). This high-resolution profiling allowed detection of rare populations present at frequencies as low as 0.05% such as Thetis III et IV in adult mice. Using this dataset, we quantified the relative abundance of each cluster in wild-type mouse MLNs. In two-week-old neonates, ILCs are the most abundant APC population, followed by cDCs, which accounted for 45.6% of total APCs (**Figure 1D**). In contrast, adult MLNs were dominated by cDCs, representing 81.5% of total APCs. Thetis cells were relatively scarce and constituted only 5.9% in neonates and less than 2.4% in adults. Additionally, a small proportion of monocyte-derived dendritic cells are also found in both neonatal and adult MLNs (**Figure 1E**).

In contrast, the composition of *Itgb8*-expressing APCs differed markedly. Itgb8(Tomato)^+^ APCs are predominantly migratory cDCs, accounting for 83.7% of Itgb8(Tomato)^+^ APCs in neonates and up to 94.5% in adults (**Figure 1F**). Within migratory cDCs, migratory cDC1 are especially enriched, representing 80% of Itgb8(Tomato)^+^ APC in adults. In neonates, Itgb8(Tomato)^+^ APCs also included 10.6% Thetis cells, with a relative enrichment of Thetis III and IV compared to total CD45^+^Lin^-^MHCII+ sample, and 3.7% ILCs. In adults, Thetis cells and ILCs accounted for less than 3% and 1% of Itgb8(Tomato)^+^ APC, respectively (**Figure 1G**).

To independently validate these observations, we performed flow cytometry analysis on MLN APC subsets isolated from *Itgb8^tdTomato^*reporter mice. Subsets were defined according to the gating strategy outlined in **Supplemental Figure S4A**. Among the populations analyzed, migratory cDC1 exhibited the highest proportion of Itgb8(Tomato)-expressing cells, with 55.6% in adults and 25.0% in neonates (**Figure 1H** and **Supplemental Figure S4B-C**). Migratory cDC2, while expressing Itgb8(Tomato) at lower frequencies, showed 6.1% expression in adults and 5.8% in neonates. Notably, Thetis cells (subsets II, III, and IV) also expressed β8, with 9.4-16.2% of cells being Itgb8(Tomato)^+^ in neonates and 6.9-9.5% in adults (**Supplemental Figure S4B-C**). However, because Thetis cells represent an extremely small fraction of total CD45^+^ cells within the MLNs, migratory cDC1 and cDC2 constitute the predominant *Itgb8*-expressing APC populations both in neonatal and adult mice, compared to Thetis cells (**Figure 4I**, top). In neonate, Itgb8(Tomato)^+^ migratory cDC1 were markedly more abundant that Thetis cells, with their frequency being 40.5 (±9) and 13.1 (±2.6) times higher than that of Itgb8(Tomato)^+^ Thetis III and IV, respectively. In adults, this difference was even more pronounced. Itgb8(Tomato)^+^ migratory cDC1 were 353 (±116.3) and 341.8 (±101.9) times more abundant that Itgb8(Tomato)^+^ Thetis III and IV, respectively (**Figure 4I**, bottom).

In conclusion, while a small fraction of RORγt⁺ APCs, particularly Thetis cells III and IV, express β8 integrin, migratory RORγt^-^ cDCs, and in particular migratory cDC1, represent the predominant β8-expressing APC population in the MLNs of both neonates and adults, supporting their potential functional role in TGF-β-mediated immune regulation.

### Generation of migratory cDC1 and cDC2-restricted β8 integrin gene inactivation models

The function of αvβ8 integrin in regulating intestinal T cell responses *in vivo* has largely been investigated using *Itgb8* conditional knockout mouse driven by *Cd11c^Cre^* (*Itgb8^ΔCd^*^11c^)^14,16,28^. While CD11c is widely considered a canonical DC marker, its expression is not exclusive to DCs, as it is also expressed by other immune cell populations, including RORγt^+^ APCs^29,30,37^ (**Figure 1C**). While *Cd11c^Cre^* efficiently targets cDCs, its activity in other MLN APC subsets complicates the interpretation of prior *in vivo* data^40^.

To specifically assess the role of *Itgb8*-expressing cDCs in pTreg induction, and to dissect the respective contributions of cDC1 and cDC2, we generated two conditional knockout models targeting *Itgb8* in each cDC lineages. For cDC1, we used the *Xcr1^Cre^* mouse model, which offer highly efficient and specific targeting across several tissues^41^. cDC2 represents a more heterogenous population, which has recently been further subdivided into cDC2A and cDC2B^21^. There is no single Cre model that comprehensively targets all cDC2. Here, we used the *huCD207^Cre^* model, which specifically targets CD103^+^CD11b^+^ cDC2 in the gut lamina propria and MLNs, while sparing other cDC subsets^26^.

To determine whether these Cre models also target other *Itgb8*-expressing APCs, such as ILC3s or Thetis cells, we performed scRNA-seq on CD45^+^Lin^-^MHCII^+^eYFP^+^ cells (‘eYFP^+^ APCs’) isolated from the MLNs of 2-week-old neonates and adult *Xcr1^cre^Rosa2C^lsl-eYFP^* and *huCD207^cre^Rosa2C^lsl-eYFP^*fate-mapping mice (**Figure 2A-D** and **Supplemental Figure S5**). As these samples were integrated with the previous samples (**Figure 1**), we projected eYFP^+^ cells onto the full APC landscape (**Figure 2 A, C**) and determined Cre activity across transcriptionally defined subsets.

**Figure 2.**
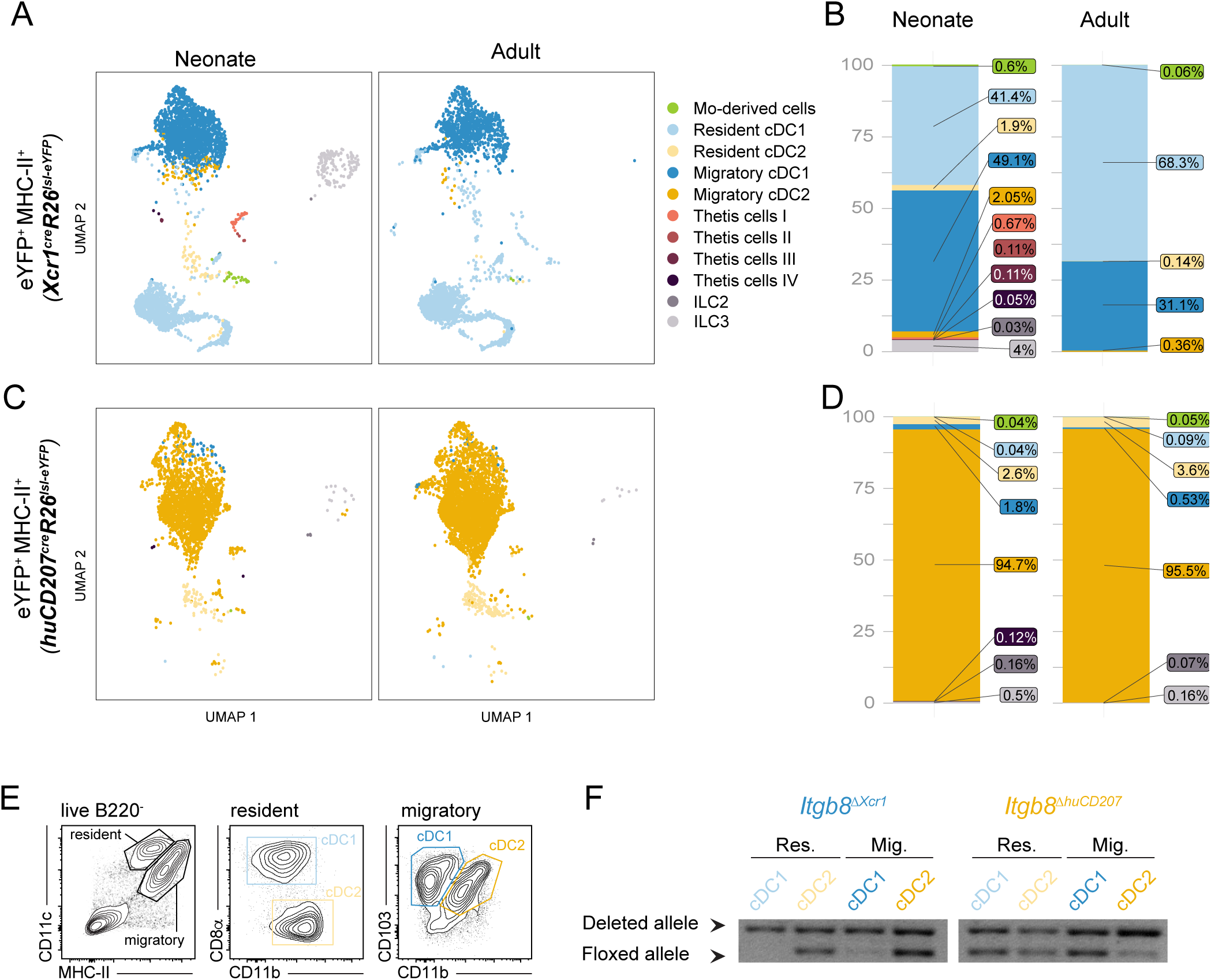
*Itgb8^ΔXcr^*^1^ and *Itgb8^ΔhuCD^*^207^ mice are selective and robust models for dissecting the role of β8 integrin in cDC1 and migratory cDC2. **A-D.** eYFP^+^ APCs (Lin^−^MHCII^+^) from the MLNs of 2-week-old neonatal and adult *Xcr1^cre^Rosa2C^lsl-eYFP^*(**A-B**) and *huCD207^cre^Rosa2C^lsl-eYFP^*(**C-D**) fate-mapping mice were sorted for scRNA-seq. **A-B**. UMAP plot (**A**) and stacked bar plots (**B**) comparing the proportion of each cluster in eYFP^+^ APCs from neonatal (left, 3,750 cells) and adult (right, 3,583 cells) *Xcr1^cre^Rosa2C^lsl-eYFP^*mice. **C-D**. UMAP plot (**C**) and stacked bar plots (**D**) comparing the proportion of each cluster in eYFP^+^ APCs from neonatal (left, 2,450 cells) and adult (right, 4,348 cells) *huCD207^cre^Rosa2C^lsl-eYFP^* mice. **E-F**. (**E**) Gating strategy and (**F**) PCR across exon 4 of *Itgb8* gene in DNA from cDC subsets sorted from MLNs of *Itgb8^ΔXcr^*^1^ and *Itgb8^ΔhuCD^*^207^ mice carrying one floxed and one deleted allele of *Itgb8*. The floxed allele is lost in resident and migratory cDC1 in mice carrying *Xcr1^Cre^* (*Itgb8^ΔXcr^*^1^, left) and only in migratory cDC2 in mice carrying *huCD207^Cre^* (*Itgb8^ΔhuCD^*^207^).

In *Xcr1^cre^Rosa2C^lsl-eYFP^* mice, cDC1 represent 90.6% of the targeted eYFP^+^ APCs in neonates and 98.8% in adults, encompassing both migratory and resident subsets (**Figure 2A-B**). A minor fraction of targeted cells in neonates comprises cDC2 (3.9%), moDCs (0.6%), Thetis cells (0.9%) and ILCs (4.0%). In contrast, *huCD207^cre^Rosa2C^lsl-eYFP^* fate-mapping revealed that 97.9% (neonates) to 99.2% (adults) of eYFP^+^ are cDC2, with a predominant migratory cDC2 signature and less that 0.7% (neonate) and 0.2% (adult) eYFP^+^ non-DCs (**Figure 2C-D**). Importantly, no eYFP^+^ Thetis cells in adult and less than 0.25% eYFP^+^ ILCs were detected in adult MLNs in either Cre model. These findings confirms that in adult mice, *Xcr1^cre^* or *huCD207^cre^* selectively target β8-expressing cDC1 and migratory cDC2, respectively. Although low-level off-target activity is observed in neonates, particularly in *Xcr1^cre^*mice, this is minimal to absent in adults.

Having demonstrated the specificity of the targeting system, *Xcr1^Cre^Itgb8^ff/-^*(*Itgb8^ΔXcr^*^1^) and *huCD207^Cre^Itgb8^ff/-^*(*Itgb8^ΔhuCD^*^207^) mice were further generated to specifically knockout β8 integrin in cDC1 and migratory cDC2, respectively. Genomic PCR on sorted MLN cDC subsets confirmed specific and significant deletion of *Itgb8* in the intended populations (**Figure 2 E-F**). In *Itgb8^ΔXcr^*^1^ mice, deletion is specific to both migratory and resident cDC1; in *Itgb8^ΔhuCD^*^207^ mice, deletion is restricted to migratory cDC2, with no deletion in resident cDC2, consistent with the known expression pattern of *huCD207*^42^ (**Figure 2E-F**). Of note, *huCD207^Cre^*-mediated recombination is not fully penetrant, and a small proportion of migratory cDC2 retained the floxed allele. Nonetheless, the specificity of these models for β8 deletion in their respective DC subsets is high.

To assess whether targeted deletion of β8 in cDC1 or migratory cDC2 alters the broader composition of MLN APCs, we quantified the proportion of cDCs, ILCs and Thetis cell subsets in *Itgb8^ΔXcr^*^1^ and *Itgb8^ΔhuCD^*^207^ mice. We observed no major differences in the frequencies of these populations in single (*Itgb8^ΔXcr^*^1^ and *Itgb8^ΔhuCD^*^207^) or double knockout (*Itgb8^ΔXcr^*^1^*^:huCD^*^207^) mice compared to littermate controls, indicating that β8 deletion in cDC1 or migratory cDC2 does not significantly impact the development or maintenance of other MLN APC subpopulations (**Supplemental Figure S6**).

In conclusion, *Itgb8^ΔXcr^*^1^ and *Itgb8^ΔhuCD^*^207^ mice are selective and robust models for dissecting the role of β8 integrin in cDC1 and migratory cDC2, respectively, and provide a powerful *in vivo* system for studying the contribution of these subsets to pTreg generation and intestinal immune homeostasis.

### cDC1 and cDC2 promote pTreg differentiation and intestinal immune tolerance in an αvβ8-dependent manner

*In vivo* studies using *CD11c^Cre^*-driven conditional deletion of β8 integrin have shown that *Itgb8^ΔCd^*^11c^ mice spontaneously develop severe colitis, associated with a two-fold reduction in the proportion of total FoxP3^+^ T cells in the colon^16^. FoxP3^+^ T cells comprise both thymus-derived (tTreg) and pTreg, the latter marked by expression of RORγt^4^. Here, we first refine that initial observation by demonstrating that *Cd11c^Cre^*-mediated β8 integrin deletion leads to a selective loss of FoxP3^+^RORγt^+^ pTreg in both the MLNs (7.4-fold reduction) and colonic lamina propria (CLP, 37.0-fold reduction) of *CD11c^cre^Itgb8^ff/-^ (Itgb8^ΔCd^*^11c^) mice compared to littermate controls (**Figure 3A-B**, top panel), while there is an increase in MLNs FoxP3^+^RORgt^-^ Tregs, which predominantly consist of Neuropilin1^+^ thymus-derived natural Treg ^43,44^ (**Supplemental Figure S7C-D**).

**Figure 3.**
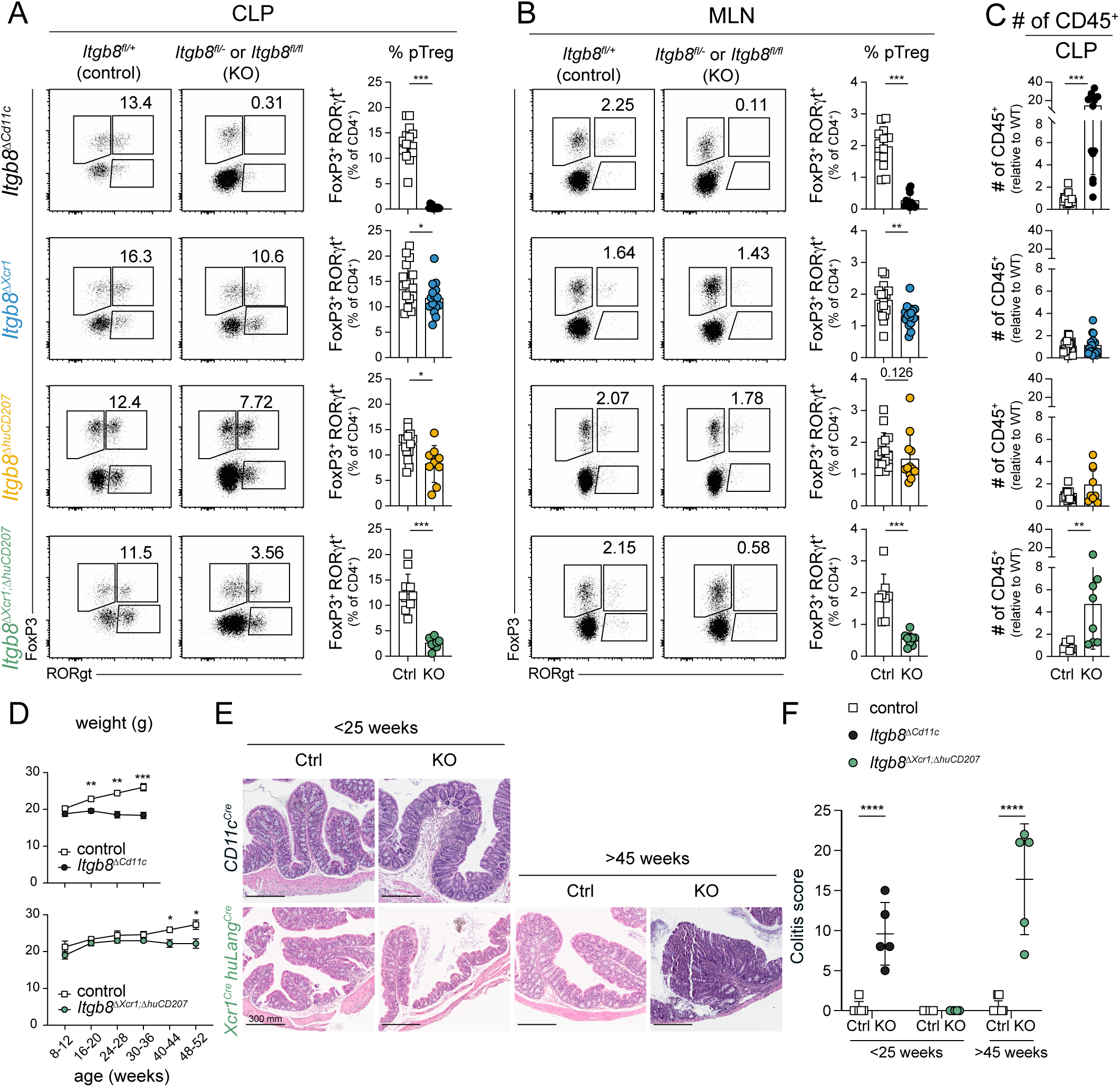
cDC1 and cDC2 promote pTreg differentiation and intestinal immune tolerance in an αvβ8-dependent manner. **A-B**. Flow cytometry of RORγt and FoxP3-expressing CD4^+^ T cell subsets (left) and summary graphs (right) for frequencies of pTreg (FoxP3^+^RORγt^+^) cells in colonic lamina propria (CLP) (**A**) and MLNs (**B**) of *Itgb8^ΔCd^*^11c^ (black circles), *Itgb8^ΔXcr^*^1^ (blue circles), *Itgb8^ΔhuCD^*^207^ (yellow circles) and *Itgb8^ΔXcr^*^1^*^;ΔhuCD^*^207^ DKO (green circles), compared to littermate control mice (open squares) (n≥9 per group). **C**. Quantification of total CD45^+^ cells in the CLP of *Itgb8^ΔCd^*^11c^ (black circles), *Itgb8^ΔXcr^*^1^ (blue circles), *Itgb8^ΔhuCD^*^207^ (yellow circles) and *Itgb8^ΔXcr^*^1^*^;ΔhuCD^*^207^ DKO (green circles), compared to littermate control mice (open squares) (n≥7 per group). **D**. Weight loss in littermate controls (open squares), and *Itgb8^ΔCd^*^11c^ (top, black circles, n=6) or *Itgb8^ΔXcr^*^1^*^;ΔhuCD^*^207^ DKO (bottom, green circles, n=5) female mice. **E**. Hematoxylin- and eosin-stained sections of colons from young (>25-week-old, left) and older (>45-week-old, right) *Itgb8^ΔCd^*^11c^ (top) and *Itgb8^ΔXcr^*^1^*^;ΔhuCD^*^207^ DKO (bottom) compared to littermate controls. **F**. Colitis score according to the Mouse Colitis Histology Index [MCHI] of *Itgb8^ΔCd^*^11c^ (black circles) and *Itgb8^ΔXcr^*^1^*^;ΔhuCD^*^207^ DKO (green circles), compared to littermate control mice (open squares); n=5 per group. *P < 0.05; **P < 0.01; ***P < 0.001; ****P < 0.0001.

To specifically dissect the contribution of β8-expressing cDCs, we analyzed *Itgb8^ΔXcr^*^1^ and *Itgb8^ΔhuCD^*^207^ mice, in which β8 is selectively deleted in cDC1 and migratory cDC2, respectively. We observed a modest but significant reduction of RORγt^+^FoxP3^+^ pTreg in both the MLNs and CLP of *Itgb8^ΔXcr^*^1^ and *Itgb8^ΔhuCD^*^207^ compared to littermate controls, with 1.2- and 1.5-fold (CLP) and 1.4- and 1.2-fold reduction (MLNs), respectively (**Figure 3A-B** and **Supplemental Figure S7A**). The relatively small effect observed in *Itgb8^ΔhuCD^*^207^ might be attributable to the partial deletion at the *Itgb8^ffox^*locus in this model (**Figure 2F**). Overall, the moderate decrease in pTreg in *Itgb8^ΔXcr^*^1^ and *Itgb8^ΔhuCD^*^207^ mice relative to that seen in *Itgb8^ΔCd^*^11c^ mice suggests either that cDC1 and migratory cDC2 make complementary contributions to pTreg induction, or that a non-cDC population, such as RORγt^+^ APC, is primarily responsible for pTreg induction. To distinguish between these possibilities, we generated mice in which both cDC1 and intestinal cDC2 are β8-deficient. These *Itgb8^ΔXcr^*^1^*^;ΔhuCD^*^207^ double knockout (DKO) mice showed a profound loss of pTreg in both MLNs (4.9-fold reduction) and CLP (3.5-fold reduction) (**Figure 3A-B** bottom panel, and **Supplemental Figure S7C**), exceeding the combined effect observed in single-deficient strains, suggesting a synergistic requirement of αvβ8 by cDC1 and cDC2.

As previously reported, loss of either αv or β8 integrin expression in CD11c-expressing APCs leads to a specific loss of the FoxP3^-^RORgt^+^ Th17 cells in the CLP and MLNs^45,46^ (**Supplemental Figure S7B**). Interestingly, our *Itgb8^ΔXcr^*^1^*^;ΔhuCD^*^207^ DKO model recapitulates this phenotype (**Supplemental Figure S7B**), albeit at a lesser extent, suggesting that additional APC subsets may contribute to β8-dependent Th17 cell differentiation. Under steady state conditions, this function appears to be primarily mediated by cDC2, as indicated by the slight reduction in Th17 cells observed in *Itgb8^ΔhuCD^*^207^ but not *Itgb8^ΔXcr^*^1^ mice. This finding is in line with the established role of cDC2 in promoting Th17 responses^24–26^.

In *Itgb8^ΔXcr^*^1^*^;ΔhuCD^*^207^ DKO mice, the profound reduction in pTreg was accompanied by a significant increase in immune cell numbers in the CLP, suggesting heightened immune activation or inflammation compared to single knockouts (**Figure 3C**). Notably, pTreg in the CLP still accounted for ∼2.5% of total T cells in *Itgb8^ΔXcr^*^1^*^;ΔhuCD^*^207^ DKO, compared to just 0.35% in *Itgb8^ΔCd^*^11c^ mice (**Figure 3A**), potentially reflecting residual contributions from non-deleted migratory cDC2 or β8-expressing RORγt^+^ APCs, such as Thetis cells.

Furthermore, contrary to *Itgb8^ΔCd^*^11c^ mice, which developed early spontaneous colitis, *Itgb8^ΔXcr^*^1^*^;ΔhuCD^*^207^ DKO mice appeared phenotypically normal during early life and only developed signs of progressive wasting disease and significant weight loss by 9 months of age (**Figure 3D**). Histological analysis of the proximal colon of *Itgb8^ΔXcr^*^1^*^;ΔhuCD^*^207^ DKO mice revealed features consistent with chronic colitis, including inflammatory infiltrates, crypt destruction, loss of goblet cell differentiation and sometimes mucosal ulceration, as confirmed by blinded pathology scoring (**Figure 3E-F**).

To assess whether the reduction in pTreg observed in DC subset-specific *Itgb8*-deficient mice also reflects impaired differentiation of food antigen-specific pTreg, we performed a food antigen-specific Treg induction assay. Naïve CD62L^+^FoxP3^-^CD4^+^ OT-II T cells were transferred into *Itgb8*-deficient or control mice, followed by oral administration of ovalbumin (OVA). Six days post-transfer, the frequency of FoxP3^+^ T cells among OVA-specific OT-II cells was only modestly reduced in *Itgb8^ΔXcr^*^1^ and *Itgb8^ΔXcr^*^1^*^;ΔhuCD^*^207^ DKO mice compared to controls, whereas no reduction was observed in *Itgb8^ΔhuCD^*^207^ mice (**Supplementary Figure 7E**). Importantly, total OT-II T cell recovery was comparable across all *Itgb8*-deficient and control groups (**Supplementary Figure 7E**). These results indicate that cDC1 contribute to the differentiation of food antigen-specific pTreg, but that this contribution is limited. This conclusion aligns with recent findings demonstrating that RORγt^+^ APCs are essential for food antigen-specific pTreg induction and the maintenance of oral tolerance^38,39,47–49^, suggesting that cDCs play only a minor role in this pathway, distinct from the more prominent one we identified in sustaining tolerance to the microbiota (**Figure 3D-F**).

### *ITGB8* is preferentially expressed by migratory cDCs in human MLNs

Previous studies have shown that in the human intestinal lamina propria, integrin αvβ8 is expressed by intestinal lamina propria and circulating CD1c^+^ cDC2, but not CD141^+^ cDC1^50^. To determine whether human DC subsets in lymphoid tissues express αvβ8 integrin, we analyzed APCs from ileum draining MLNs of 3 Crohn’s disease patients by scRNA-seq (**Supplemental Table 1**). After preprocessing the data, samples were integrated and cDCs were bioinformatically isolated. Five APC subsets were identified based on unsupervised clustering and expression of signature gene scores (**Figure 4A-B**). CCR7^+^ DC cluster expressed predominantly genes related to DC maturation and migration (**Supplemental Figure 8A**). On querying for *ITGB8* in this human dataset, we observed preferential expression in migratory CCR7^+^ cDCs. (**Figure 4C and supplemental Figure 8B**).

**Figure 4.**
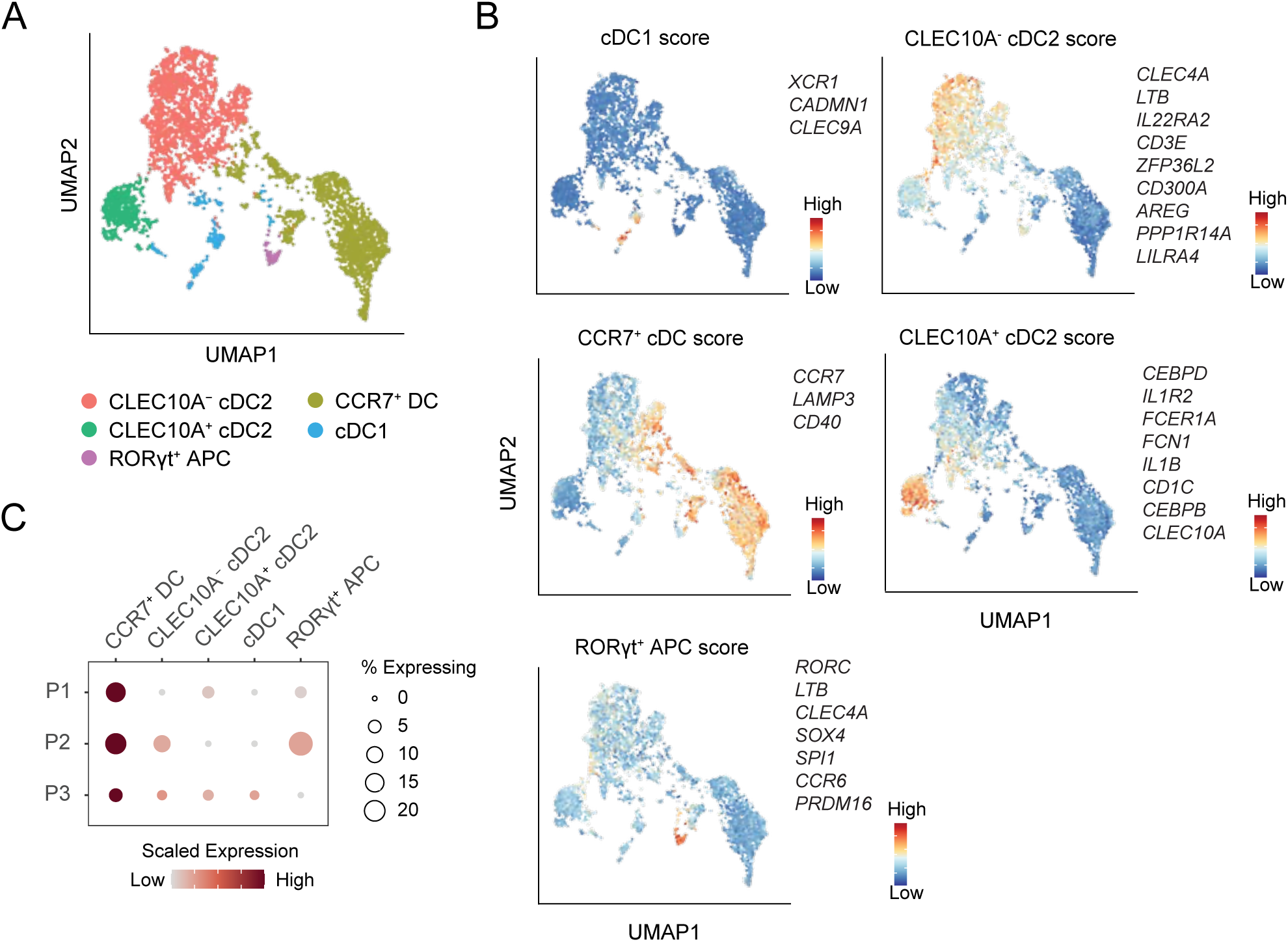
*ITGB8* is preferentially expressed by CCR7^+^ migratory DCs in human MLN. **A.** UMAP visualization of cells obtained from scRNA-seq analysis of small intestine draining lymph nodes (SI-LN) from 3 Crohn’s Disease (CD) patients, colored by cluster annotation. **B**. APC subset scores. CLEC10A^-^ cDC2 score and CLEC10A^+^ cDC2 score is from Brown *et al.*^21^. RORγt^+^ APC score is from Antonova *et al.* ^61^. **C**. Dot plot showing the expression of *ITGB8* in human MLN APC subsets.

Taken together, these results underscore the essential role of TGF-β activation by cDCs, via αvβ8 integrin, in the differentiation of intestinal pTreg. Both cDC1 and migratory cDC2 contribute to this process in a complementary, partially redundant, manner. In the absence of αvβ8 on both subsets, pTreg differentiation is severely impaired, leading to immune dysregulation and spontaneous colitis with age, emphasizing the non-redundant role of αvβ8 integrin in maintaining intestinal immune tolerance through adulthood. At last, our data showing that *ITGB8* is expressed by human MLN cDCs underscore the potential conserved nature of this mechanism between mice and humans.

## DISCUSSION

Using a murine integrin β8 (*Itgb8*) gene reporter model combined with single-cell profiling, we provide here a comprehensive characterization of *Itgb8*-expressing APCs capable of promoting pTreg differentiation through αvβ8-mediated TGF-β activation. Our findings reveal that *Itgb8* is predominantly expressed by migratory cDC1 and cDC2, in both neonatal and adult mice. Functional experiments further demonstrate that selective deletion of *Itgb8* in either cDC1 or cDC2 results in reduced intestinal pTreg frequencies, implicating both subsets in pTreg induction. Notably, spontaneous colitis arises only when β8 is concurrently deleted in both subsets, suggesting a degree of functional compensation between cDC1 and cDC2 in maintaining tolerance.

Nevertheless, the results underscore the essential, functionally specialized roles of cDCs in intestinal immune regulation, challenging recent studies that have questioned their requirement for pTreg induction^29–31^. Importantly, our genetic targeting strategy specifically disrupts *Itgb8* expression in *bona fide* cDCs, while sparing RORγt^+^ APCs. This minimizes the possibility that the observed effects are due to off-target deletion in other APC populations, such as RORγt⁺ cells, and strengthens the conclusion that cDC subsets themselves are sufficient to drive pTreg differentiation via αvβ8-mediated TGF-β activation.

Historically, migratory CD103⁺ cDCs in the MLNs have been implicated in pTreg generation, acting through TGF-β-dependent mechanisms to mediate tolerance to gut-derived antigens^10,11,51–53^. Their capacity to activate latent TGF-β through αvβ8 integrin expression has made them prime candidates for mediating peripheral tolerance^13,14^. However, CD103⁺ cDCs represent a heterogeneous population encompassing both cDC1 and cDC2, each with distinct transcriptional and functional profiles^18^. While *in vitro* studies had previously demonstrated that cDC1 are especially proficient at inducing pTregs and activating TGF-β^27,28^, the *in vivo* relevance of these observations has remained uncertain.

Most *in vivo* models assessing cDC subsets function have largely relied on the deletion of transcription factors critical for lineage specification (e.g. *Irf8^ΔCD^*^11c^ and *Batf3^-/-^* mice for cDC1, and *Irf4^ΔCD^*^11c^ for cDC2), or on ablation using subset-specific expression of suicide gene (e.g. *XCR1-DTA* for cDC1, *HuCD207-DTA* for migratory cDC2)^54^. However, these models are limited by compensatory mechanisms, such as cellular plasticity or incomplete deletion, which may obscure the contributions of residual or reprogrammed DCs. For example, *Irf8^ΔXcr^*^1^ and *Batf3*^⁻^*^/^*^⁻^ mice show cDC1 adopting cDC2-like phenotypes^55^, and similar lineage reprogramming is observed in *Batf3^-/-^* mice^56^. Consequently, prior conclusions suggesting that cDCs are dispensable for pTreg induction must be revisited considering this limitation.

These issues are highlighted by the stark contrast between cDC-deletion models and *Itgb8^ΔCD^*^11c^ mice, in which deletion of either αv or β8 integrin in all CD11c⁺ cells result in profound loss of Foxp3⁺ Tregs and spontaneous colitis^16,17^. This discrepancy led to speculation that non-cDC CD11c⁺ APCs, particularly the recently identified RORγt⁺ APCs, might be the primary drivers of pTreg induction. Indeed, three seminal studies in 2022 identified RORγt⁺ MHC-II⁺ APCs, including eTACs, ILC3s, and Thetis cells, as essential for tolerance to microbiota, with *Rorc^Cre^ × H2-Ab1^ff/ff^*mice showing marked pTreg loss upon MHC-II deletion in RORγt⁺ cells^29–31^.

However, the interpretation of these results is complicated by the potential for transient *Rorc* expression in cDCs during their development or maturation in the intestinal lamina propria. Indeed, we recently identified a population *RORC*^+^*PRDM1C*^+^ cells in the human intestinal LP, that in trajectory-based analysis appeared to lie upstream of cDC2^57^. Furthermore, fate-mapping studies have shown that a subset of murine CD11c⁺ cDCs retain a history of RORγt expression, despite no longer expressing RORγt anymore in the MLNs^11^. Thus, the *Rorc^Cre^* driver is not restricted to RORγt^+^ APCs (Thetis cells, eTACs, tolDCs and ILCs) in the MLNs, but can also target cDCs, irrespective of their current RORγt expression status. This raises the possibility that *Rorc^Cre^*-driven deletions might inadvertently affect cDCs, thereby blurring the distinction between these novel APCs and classical cDCs. Moreover, studies using *ClecSa^Cre^ × H2-Ab1^ff/ff^* mice to selectively delete MHC-II in cDCs are limited by incomplete recombination efficiency, leaving a substancial proportion of cDCs functionally intact and capable of sustaining pTreg responses^29,32,33^.

In contrast to these earlier models, our study uses refined genetic tools to selectively target β8 expression in cDC1 and cDC2 without affecting RORγt⁺ APCs, allowing a more precise evaluation of each cDC subset contribution to pTreg induction. We demonstrate that deletion of *Itgb8* in either cDC1 or cDC2 leads to a drastic reduction in pTreg, and that the combined deletion in both subsets has an additive effect, resulting in colitis in β8-DKO mice. These findings clearly establish that canonical cDCs are both sufficient to induce pTregs and essential for maintaining immune homeostasis in the gut.

In summary, our data support a model in which multiple cDC populations—cDC1, cDC2—contribute in a partially redundant yet essential manner to pTreg differentiation in adult mice. The lack of phenotype in young mice suggests that tolerance may instead be initially established by RORγt⁺ APCs during development; in particular, in the first weeks after birth as these cells are enriched in neonates (Fig.1 and ^29^). Additionally, others have shown that RORγt⁺ APCs play a role under pathological conditions^30,31^, which we have not ruled out in our study. This apparent redundancy among MLN APC populations likely reflects an evolutionary pressure to ensure robust tolerance mechanisms in the antigen-rich, dynamic intestinal environment. Importantly, despite their differences in origin and phenotype, these APC subsets converge on a shared functional mechanism: the activation of latent TGF-β via αvβ8 integrin. This pathway appears central for the induction of peripheral tolerance in the gut and may also play a role in systemic immune regulation, such as what was recently demonstrated for the induction of allograft antigen-specific Treg induction and tolerance in transplantation (Zhang et al, Nature *In press*). This integrative model offers a new perspective that helps reconcile previously conflicting findings, such as the minimal effect of cDC depletion on pTreg numbers in certain settings and underscores the complexity and temporal specificity of immune tolerance mechanisms in the gut.

Finally, we show that, in humans, *ITGB8* is expressed by MLN CCR7^+^ migratory cDCs. This finding aligns with a broader body of research highlighting the pivotal role of αvβ8 integrin in immune regulation. Notably, a 2017 Genome-Wide Association Study (GWAS) first identified *ITGB8* as a susceptible locus associated with human pathology^58^. Interestingly, the associated genetic variant in *ITGB8* correlated changes in *ITGB8* expression with development of Inflammatory Bowel Disease (IBD). In parallel, in 2018 a study in mice showed that induction of αvβ8 expression in DCs via probiotic administration conferred protection against colitis^59^. These data, combined with recent studies demonstrating the therapeutic potential of αvβ8 targeting in inflammatory disorders beyond the gut, such as encephalitis or lung fibro-inflammation^46,60^, underscore its broader clinical relevance. Further studies will be required to determine the relationship between *Itgb8*-expressing MLN DC subsets and the recently characterized RORγt⁺ APCs in the human intestinal lamina propria^38,61^. Nevertheless, given its critical immunoregulator role and shared expression in human cDCs, αvβ8 integrin emerges as a promising target for therapeutic modulation of immune tolerance. This paves the way for novel diagnostic and therapeutic strategies aimed at restoring immune homeostasis in inflammatory, autoimmune, and allergic diseases, as well as inducing graft tolerance in transplantation.

## METHODS

### Mice

All animals used in this study were on a C57BL/6 background and maintained in specific-pathogen-free facilities at Plateau de Biologie Expérimentale de la Souris (Lyon, France). Sex-matched littermate controls were used for all experiments, with both male and female mice included indiscriminately. All animal studies and procedures were conducted in compliance with European Union guidelines and were approved by the local ethical committee for animal research (CECCAPP Lyon) which is registered with the French National Ethics Committee of Animal Experimentation under no. C2EA15.

Transgenic *Cd11c*^Cre^ (B6.Cg-Tg(Itgax-cre)1-1Reiz/J) (Caton 2007 17591855) were obtained from Nathalie Bendriss-Vermare (CRCL, France), *Xcr1^Cre^* (Xcr1-IRES-iCre-2A-mTFP1 B6-Xcr1tm2Ciphe) from Bernard Malissen (CIML, France) and *HuCD207*^Cre^ (B6.Cg-CD207^tm2^.^1^(cre)^Bjec^) mice^42^ from Katharina Lahl (Lund University, Sweden), *Itgb8*^flox/flox^ (Itgb8^tm2Lfr^)^62^ obtained from Julien Marie (CRCL, France) or *Rosa2C^lsl-eYFP^* (B6.129X1-Gt(ROSA)26Sortm1(EYFP)Cos/J^63^ obtained from François-Loïc Cosset (CIRI, France); OT-II mice (B6.Cg-Tg(TcraTcrb)425Cbn/J)^64^ crossed to *Foxp3^eGFP^* (B6.Cg-Foxp3tm1.1Mal/J)^65^ mice were obtained from Marc Vocanson. *Itgb8*^tdTomato^ (Itgb8-IRES-tdTomato) were previously described^34^. When crossed to Cre-expressing lines, both *Itgb8^ff/ff^* and *Itgb8^ff/-^*were used as β8-KO mice.

### Tissue processing and isolation of immune cells

Single-cell suspensions from the spleen and MLNs were obtained by mechanical disruption through a 100 µm cell strainer in Complete RPMI buffer (RPMI-1640 supplemented with 10% FBS, 1% Pen/Strep (Gibco), 1% HEPES (Gibco) and 0.1% β-mercaptoethanol (all Gibco)). Mononuclear phagocytes from the spleen and MLNs were isolated as previously described (Boucard-Jourdin et al. 2016). Briefly, spleen and MLNs were digested in RPMI containing 1 mg/mL Collagenase IA (Sigma) and 100 µg/mL DNAse 1 (Roche). Colon lamina propria cells were isolated as previously described. Briefly, mucus and epithelial cells were removed with 1.5 µg/mL DTE (Sigma) and 10 mM EDTA solution before digestion with 12.5 µg/mL Liberase TM (Roche) and 100 µg/mL DNAse1 (Roche). Mononuclear cells were then enriched using a Percoll gradient.

### Flow cytometry

The antibodies used for flow cytometry and FACS are listed in Supplemental table 1. For flow cytometric analysis, dead cells were excluded by staining with Fixable Viability Dye eFluor780 or Fixable Viability Dye eFluor506 (Invitrogen). Cells were then stained for surface antigens in the presence of anti-CD16/32 to block binding to Fc receptors for 30 min at 4°C in PBS. For intranuclear protein analysis, cells were fixed and permeabilized with eBioscience™ Foxp3/Transcription Factor Staining Buffer Set according to manufacturer instructions. For intranuclear protein analysis of samples from *Itgb8^tdTomato^*mice, protocol was adapted as follows: following surface staining, cells were fixed with ROTI®Histofix (4% formaldehyde) for 1 h at RT then washed two times 10 min with Permeabilization buffer 1X from the eBioscience™ Foxp3 / Transcription Factor Staining Buffer Set. Intranuclear staining was performed overnight at 4°C. Data acquisition was performed using LSR-Fortessa and Cytek Aurora spectral flow cytometers while DC subsets were sorted using a BD FACSAriaII cytometer (BD Biosciences). Data analysis was performed using FlowJo software v10 (Treestar Inc).

### Histological analysis of intestinal inflammation

A short piece of colon in the proximal section was harvested and fixed in formalin 4% for 24 h. Samples were then kept at 4°C in ethanol 70% until dehydration, clearing and paraffin embedding. Cross-sections of large intestine were stained with hematoxylin and eosin and graded by a trained pathologist (T.F.) using the Mouse Colitis Histology Index [MCHI] described by Koeling and colleagues^66^, taking into account goblet cell loss, crypt density, hyperplasia and submucosal infiltrate. The total score of the MCHI ranges from 0 [no disease] to 22 [severe disease]. Reader of histological slides was blinded to the experimental groups. Images were obtained using Aperio ImageScope viewer v12.4.3.5008 software.

### PCR, RT-qPCR and Western Blot analysis of beta 8

Before FACS sorting, filtered cell suspensions were enriched for DC using an OptiPrep gradient (Sigma). MLN dendritic cells were FACS-sorted and genomic DNA was extracted using Proteinase K, as previously described^28^. Cre-mediated deletion in *Itgb8* locus was analysed by PCR on genomic DNA using the following set of primers: 5’-CCC ACT AAG ATA ACT GGC CGT ATC-3’, 5’-GAG GGG TGG GGA AAT TTT TGT ATC-3’, 5’-GTG GAT TCT ACA GGC AAG C-3’.

### Murine scRNA-sequencing

#### Cell sorting and fixation

Mesenteric lymph nodes were collected from neonate (2-week-old) and adult (14 - 40-week-old) *Xcr1^cre^Rosa2C^lsl-eYFP^*, *huCD207^cre^Rosa2C^lsl-eYFP^*, *Itgb8^tdTomato^*mice and control mice. Tissues were digested as described above and cells were labelled for magnetic column enrichment (Miltenyi LD column depletion with streptavidin microbeads and biotinylated antibodies against TCR𝛾/𝛿, TCRβ, NK1.1, B220 and GR-1. Cells were then sorted by flow cytometry for live Lin (TCR 𝛾/𝛿, TCRβ, NK1.1, B220 and GR-1)^-^, CD45^+^, MHC-II^+^, eYFP^+^ for *Xcr1^cre^Rosa2C^lsl-eYFP^*, *huCD207^cre^Rosa2C^lsl-eYFP^ mice*; live Lin^-^, CD45^+^, MHCII^+^, tdTomato^+^ for *Itgb8^tdTomato^* mice and live Lin^-^, CD45^+^, MHCII^+^ for control mice. Cells were fixed using the Evercode cell fixation V2 (ECF2001, Parse Biosciences) following manufacturer’s instructions. Briefly, cells were fixed and permeabilized, counted and frozen in the presence of DMSO 5%. All samples were then stored at -80°C until library preparation.

#### Library preparation and scRNA sequencing

Evercode’s Whole Transcriptome (WT) v2 kit (ECW02030, Parse Biosciences) was used for the preparation of single cell libraries for RNA sequencing. Eight samples containing a total of 100,000 cells to barcode were loaded in a 96-well plate provided by Parse Biosciences for 3 rounds of barcoding. The targeted number of barcoded cells were 10,000 cells for neonate and adult *Xcr1^cre^Rosa2C^lsl-eYFP^* and *Itgb8^tdTomato^*, 6,000 and 8,000 cells for neonate and adult *huCD207^cre^Rosa2C^lsl-eYFP^*, respectively and 23,000 cells for neonate and adult control B6 mice. After counting the cells, 8 separate aliquots of 12,500 cells per sublibrary were processed in parallel. For each sublibrary, cells were lysed, barcoded cDNA was amplified, fragmented, and a unique sublibrary index was added prior to sequencing. The eight sublibraries were sent to Macrogen for NovaSeqX (Illumina) sequencing and an average of 41,805 reads/cells (R1+R2) were generated.

### Murine scRNA-seq computational analysis

#### Pre-processing

The quality of sublibraries were checked using FastQC (v0.12.1). Then, we followed the alevin-fry tutorial (https://combine-lab.github.io/alevin-fry-tutorials/2022/split-seq/) to process the data. Splitp v0.2.0 (without allowed mismatch) was used to combine oligo-dT amplification barcodes from the first round of the Parse Bioscience protocol to those corresponding to random hexamers. Mouse reference genome (GRCm39) was downloaded from Ensembl (v112) and a splici index was created using the roe R package using transcripts sequences, intronic regions and Tomato/YFP fluorophore sequences. This index was used to quasi-mapping and quantification using alevin-fry [10.1038/s41592-022-01408-3] from Salmon (v1.10.3) [10.1038/nmeth.4197] and thus to obtain a cell-by-gene count matrix for each sample.

#### QC and normalization

Count matrix was then processed and analyzed using R (v4.4.1) and Seurat (v5.1.0). Preliminary quality control of each sample was performed using miQC (v1.12.0) to remove low-quality cells with a posterior cutoff of 0.75 independently for each sample. Normalization was performed using the log normalization method.

#### Dimension reduction, cell clustering, and visualization

Data from all experimental conditions were then scaled, and dimensionality reduction was performed using Principal Component Analysis (PCA), retaining the top 50 PCs. To integrate the datasets, the function IntegrateLayers with the Canonical Correlation Analysis (CCA) approach was used, with PCA as the initial dimension reduction method. Subsequent non-linear dimensionality reduction was conducted using UMAP with default parameters. Clustering was performed using the Louvain algorithm, implemented in Seurat’s FindClusters function, with a resolution of 0.04. Before annotation, doublet predictions were made using DoubletFinder (v1.18.0) with the top 25 principal components (PCs) assuming a 10% doublet formation rate to remove doublets. For the initial analysis presented in the results, we focused on total and Itgb8(Tomato)^+^ APCs (CD45^+^Lin^−^MHCII^+^) isolated from healthy neonate (2-week-old) and adult (age range) Itgb8-IRES-tdTomato (Itgb8tdTomato) reporter and control B6 mice. YFP^+^ subsets, while included during integration and clustering, are introduced in subsequent sections to address their specific transcriptional profiles.

#### Clusters annotation

Cell type identities per cluster were assigned based on a combination of canonical marker genes, specific cluster marker genes, and comparison with published datasets. Cluster-specific markers were identified using Seurat’s FindAllMarkers function with a minimum expression threshold of 60% (min.pct = 0.6). The annotation of ILCs and Thetis cell subpopulations was guided by projecting the dataset from Lyu *et al.* (GSE184175)^31^ and Akagbosu *et al.* (GSE205066)^29^, respectively, onto our dataset using Seurat’s FindTransferAnchors and TransferData functions. Finally, two minor contaminant clusters (B cells and glial cells) were identified and excluded from downstream analysis.

### Human lymph node collection, cell isolation and scRNA-seq

Ileal draining mesenteric lymph nodes were collected from terminal ileum resections from 3 Crohn’s disease patients (**Supplemental Table 1**) undergoing surgery for disease relief after informed consent with ethical approval from the Scientific Ethics Committee of the Copenhagen Capital Region, Denmark (H-20054066). Lymph nodes were cleaned from surrounding mesenteric tissue, cut into smaller pieces and digested for 45 min in R5 (RPMI/5% FCS/1% penicillin and streptomycin) containing DNase 1 (30 μg/ml) and collagenase D (5 mg/ml) at 37 °C under gentle shaking (370 rpm). Resulting cell suspensions were passed through 100 μm filter and washed with R5.

Immune cells were isolated by fluorescently activated cell sorting and subjected to single cell RNA sequencing using the Chromium Next GEM Single Cell 5’ Reagent Kits v2 (Dual Index) with Feature Barcode technology for Cell Surface Protein & Immune Receptor Mapping (10x Genomics) following the manufacturer’s instructions. Quality and quantity of final libraries were measured using the Agilent 2100 Bioanalyzer equipped with High Sensitivity DNA chip (Agilent). Sequencing was carried out at the SNPCSEQ Technology Platform, Sweden or at the Core Facility for Flow Cytometry and Single Cell Analysis, Faculty of Health and Medical Sciences, University of Copenhagen using Illumina NovaSeq systems (300 cycles) aiming for a minimum of 30,000 read pairs/cell.

### Human scRNAseq computational analysis

Sequencing data was pre-processed and aligned with 10x Genomics Cell Ranger (version 8.0.0). Each sample was read into a Seurat (version 5.2.1)^67^ object in R (version 4.3.1) and processed by removing cells with exceptionally low or high UMI, gene counts or mitochondrial gene content according to the best practice^68^. Doublets were predicted and removed using scDblFinder (version 1.16.0)^69^. After log-normalization of RNA levels for individual samples, cell cycle gene modules were calculated using Seurat CellCycleScoring function, variable genes were identified, and samples were integrated with harmony^70^. Mononuclear phagocytes were bioinformatically isolated, and plasmacytoid dendritic cells (*IL3RA*, *CLEC4C*, *GZMB*) as well as monocytes-macrophages (*S100A8*, *CD14*, *C1QA*, *CSF1R*) were removed before analysing *ITGB8* expression levels.

### Statistical analysis

Detailed information on sample sizes, number of biological replicates, statistical tests used, is provided in the Methods section and figure legends. Statistical significance was evaluated as indicated in each figure legend using GraphPad Prism 10 software (GraphPad).

### Graphical illustrations

Graphical elements used to illustrate experimental design schemes were created with BioRender.com

## Data and code availability

The raw and processed murine single-cell RNA-seq data have been deposited in ArrayExpress under accession number E-MTAB-15839 and will be released upon publication. The processed data for human single-cell RNA-seq is shared upon request from the authors. The data will be made publicly available in ArrayExpress at the time of publication. Scripts used to reproduce the murine scRNA-seq analyses are available at: https://gitbio.ens-lyon.fr/ciri/nopab/delphine-bv/this-brichart-vernos-2025 and scripts for the human scRNA-seq analyses can be accessed at: https://github.com/venlavaan/This_Brichart-Vernos_scRNAseq_Figure_4_and_S8.

## Acknowledgements

We acknowledge the contribution of SFR Biosciences (Universite Claude Bernard Lyon 1, CNRS UAR3444, Inserm US8, ENS de Lyon) facilities, in particular the Plateau de Biologie Expérimentale de la Souris (PBES), and the flow cytometry facility (AniRA-Cytométrie). Elements of cartoons in Figure 1A or 4A were created with BioRender.com. This work was supported by the Agence Nationale de la Recherche (ANR-20-CE15-0015) (H.P.), a PhD fellowship from the French Ministry of Higher Education (to S.T.) and grants from the Swedish Medical Research Council (2022-00591) and the Novo Nordisk Foundation (grant number NNF22OC0071681) to W.W.A. We gratefully acknowledge support of the Institut Français de Bioinformatique (IFB, CNRS UMS 3601) for the computing resources. The IFB is funded by the Programme d’Investissements d’Avenir (PIA), grant from the Agence Nationale de la Recherche, number ANR-11-INBS-0013.

## Author Contributions

S.T., D.B-V. and H.P. conceived and designed the study; S.T., D.B.-V., V.A.V. , V.B., U.M.M., L.D. and H.P. performed experiments and analyzed data; D.B-V., V.A.V. and C.R. performed computational analyses; T.F. performed imaging analyses of mouse tissue; U.M.M., S.L.N. and S. G-D. provided access to key reagents and human samples. O.T. and W.W.A. provided advice on conceptualization and experimental design, and discussed data; S.T., D.B-V. and H.P. wrote the manuscript with input from the other authors; H.P. supervised the study and acquired funding; all authors read and approved the manuscript.

---

**Supplemental Figure S1.**
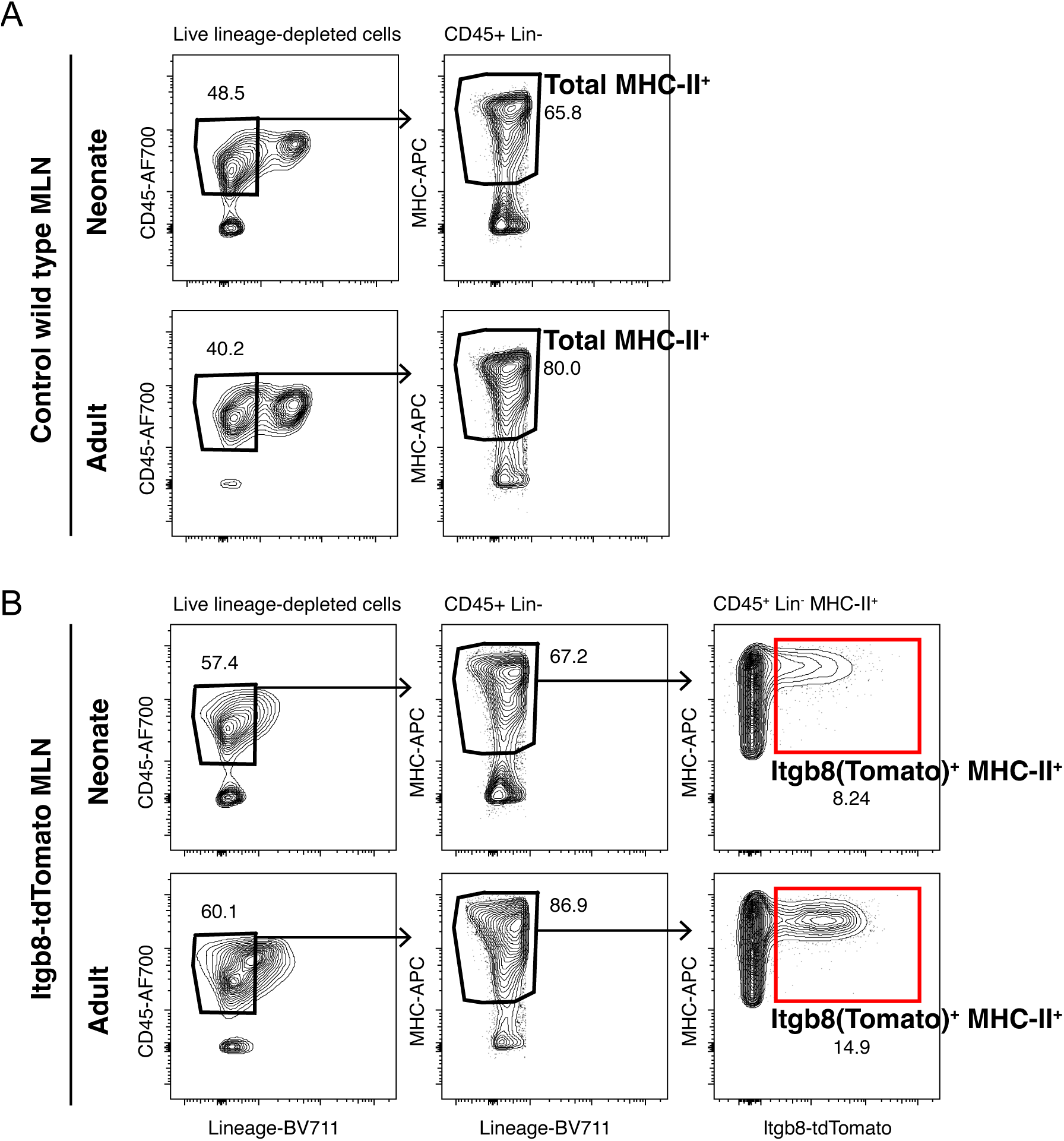
Gating strategy for scRNA-seq of total and Itgb8(Tomato)^+^ APC. **A-B**. Gating strategy used to sort total (**A**) and Itgb8(Tomato)^+^ Lin^−^MHCII^+^ (**B**) cells from the MLNs of 2-week-old neonatal (**top**) or adult (**bottom**) of control (**A**) or Itgb8-IRES-tdTomato (*Itgb8^TdTomato^*) reporter mice (**B**).

**Supplemental Figure S2.**
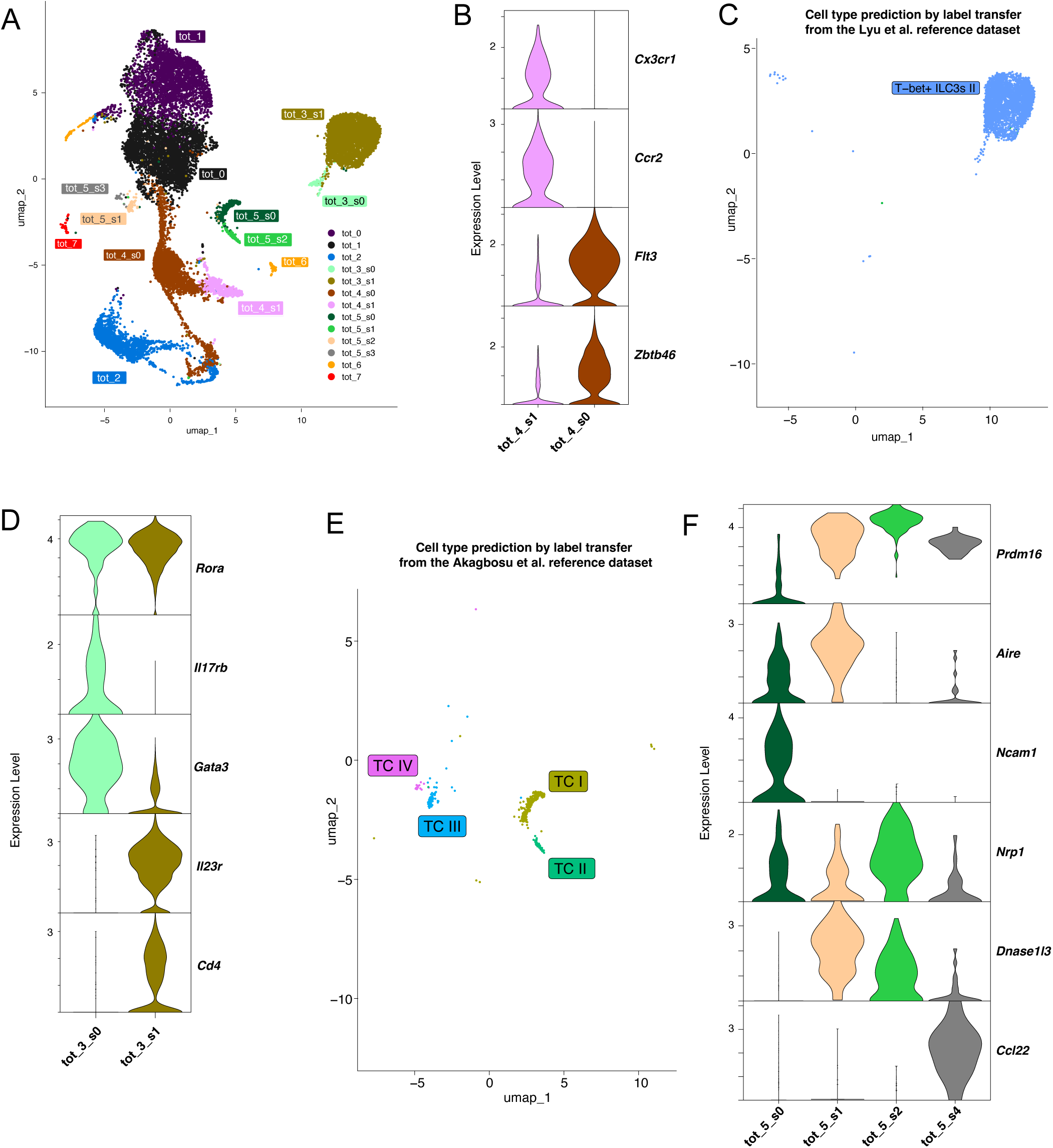
Annotation of specific APC subsets in the MLN APC atlas. **A-B.** UMAP visualization of MLN APC clusters prior to annotation, before (**A**) and after (**B**) subclustering. **C**. Violin plots showing the expression of canonical monocyte-derived cell and dendritic cell marker genes in subclusters cluster_4_s0 and cluster_4_s1. **D**. UMAP of the cluster_3 subset, with predicted cell type annotations inferred by reference mapping to the dataset from Lyu et al.^31^. **E**. Violin plots showing the expression of canonical ILC, ILC2 and ILC3 marker genes in subclusters cluster_3_s0 and cluster_3_s1, used to distinguish ILC subtypes. **F**. UMAP of the cluster_5 subset annotated by reference mapping to the dataset from Akagbosu et al.^29^. **G**. Violin plots showing the expression of Thetis cells marker in subclusters cluster_5_s0, cluster_5_s1, cluster_5_s2 and cluster_5_s3, used to identify Thetis I-IV subsets.

**Supplemental Figure S3.**
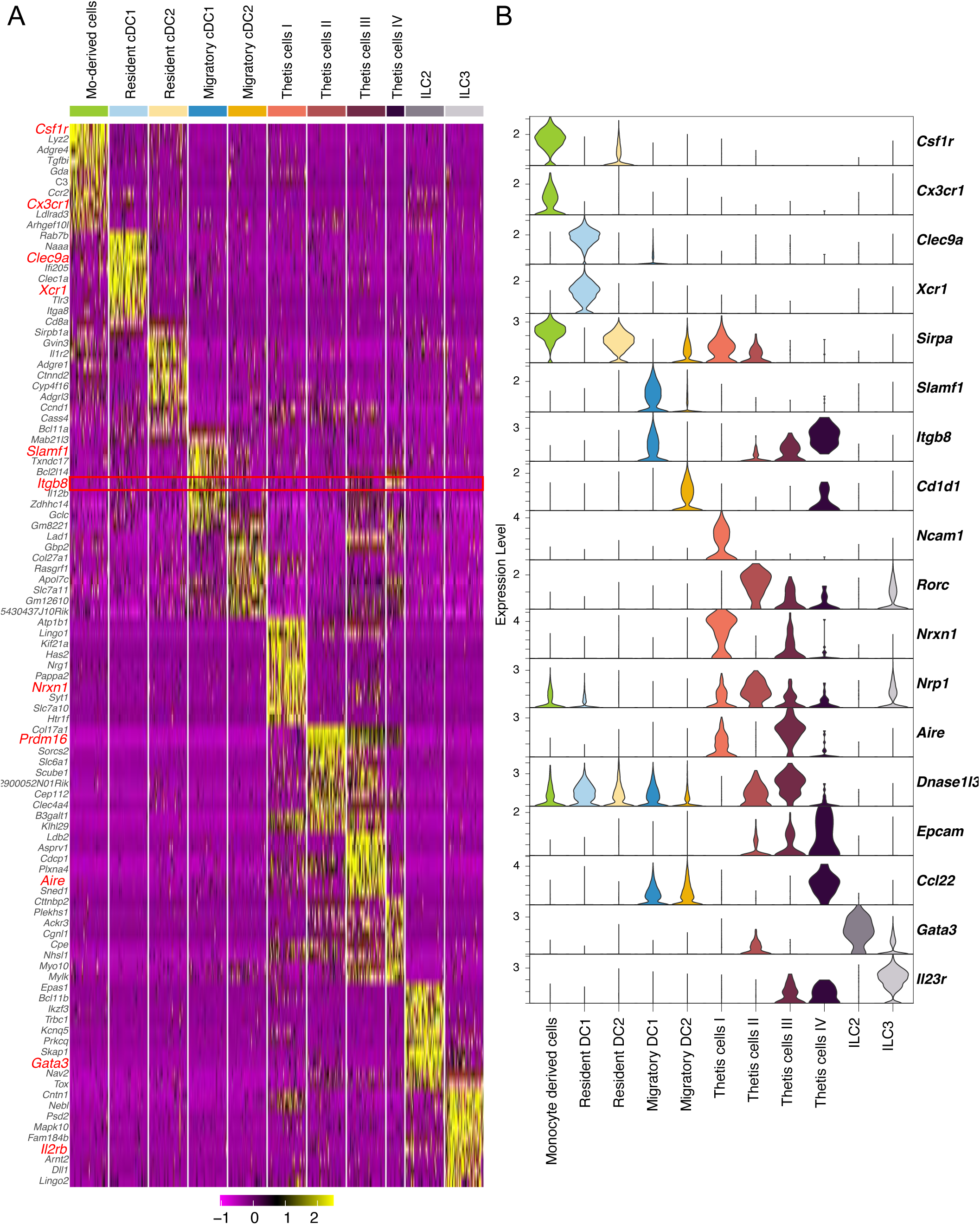
Transcriptional signatures of MLN APC subsets. **A.** Heatmap showing the top10 differentially expressed genes (DEGs) per cluster, computed from cells expressing ≥60% of the gene with a log_2_ fold-change > 1 when compared to all other clusters. Genes were selected on highest average log_2_FC across clusters. For visualization, 50 cells per clusters were downsampled. The *Itgb8* gene is outlined in red; additional canonical genes used for cluster annotation are also highlighted in red. **B**. Violin plots displaying expression of representative marker genes across APC subsets defined in the integrated MLN atlas.

**Supplemental Figure S4.**
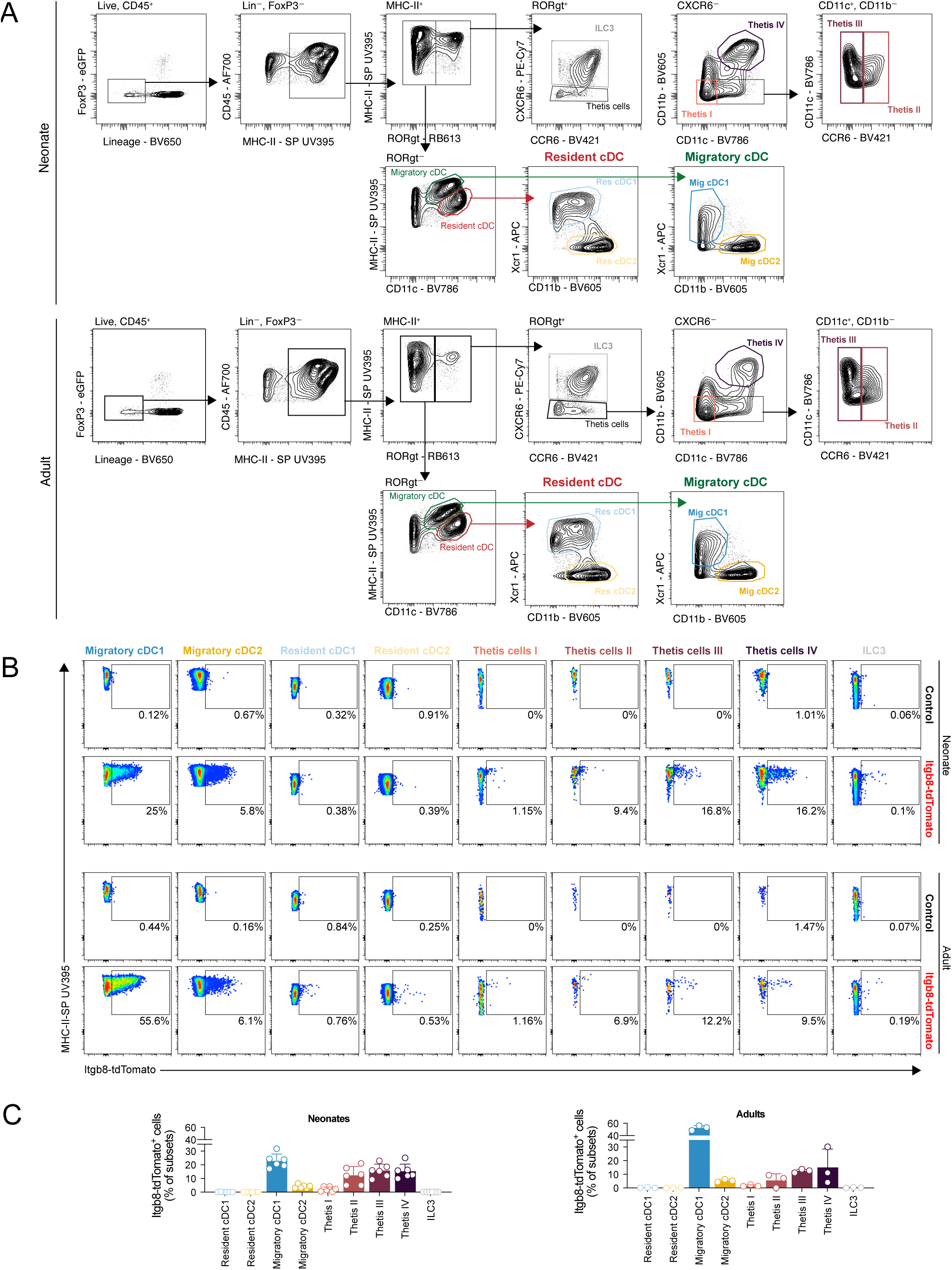
Flow cytometric validation of β8 expression across MLN APC subsets. **A**. Gating strategy used to identify APC subsets from MLNs of neonatal (top) and adult (bottom) mice. All plots represent concatenated samples from *Itgb8^tdTomato^* reporter mice (n=6 adults; n=3 neonates). **B**. Representative flow cytometry plots showing β8(tdTomato) expression in individual APC subsets from reporter mice, compared to littermate control mice lacking the tdTomato reporter. Percentages indicate the proportion of tdTomato⁺ cells within each gated subset. **C**. Proportion of Itgb8(Tomato)^+^ cells for each APC subset.

**Supplemental Figure S5.**
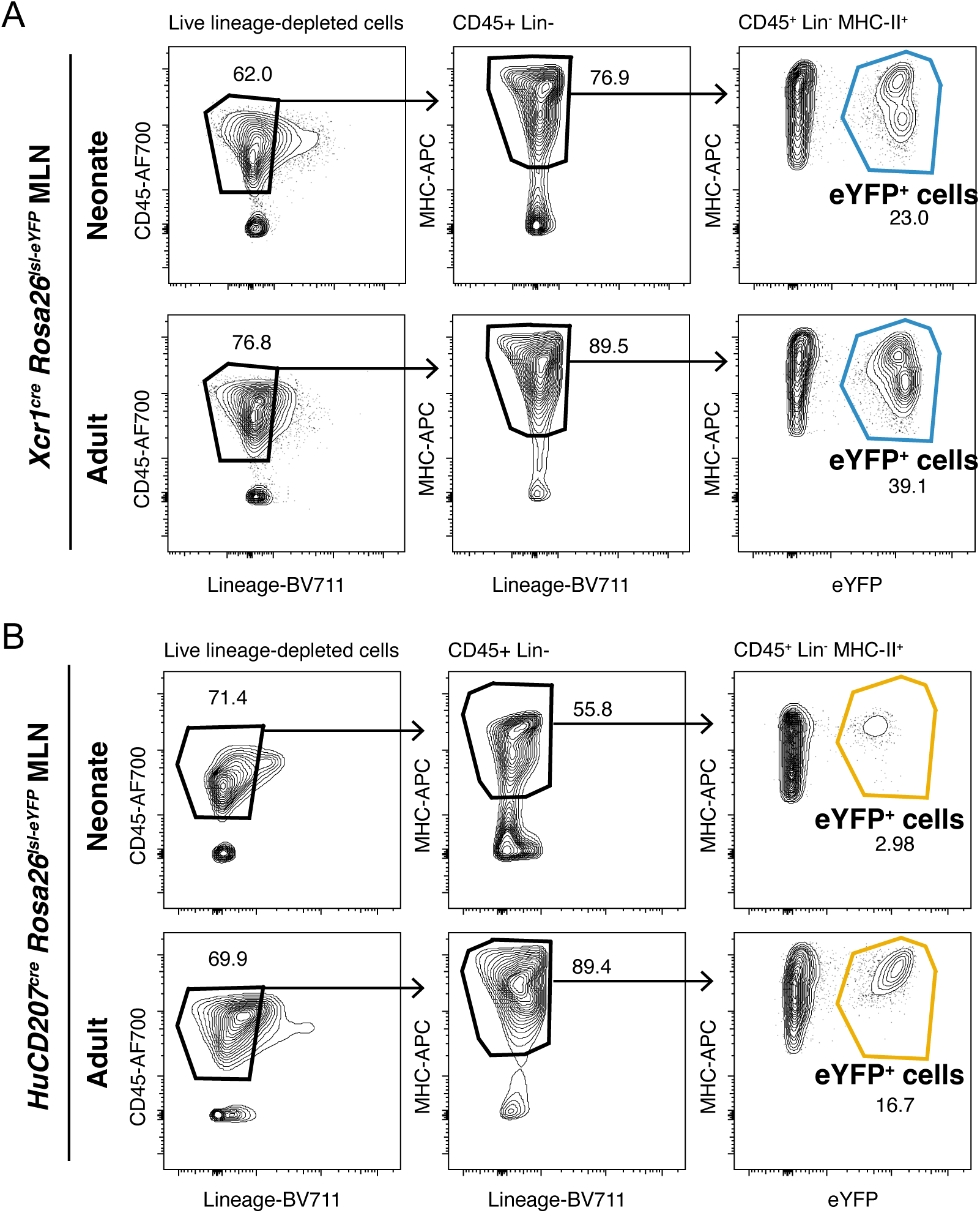
Gating strategy for scRNA-seq of eYFP^+^ APCs from fate-mapping models. **A-B**. Gating strategy used to sort CD45^+^Lin^-^MHCII^+^eYFP^+^ cells from the MLNs of 2-week-old neonatal (**top**) or adult (**bottom**) of *Xcr1^cre^Rosa2C^lsl-eYFP^* (**A**) or *huCD207^cre^Rosa2C^lsl-eYFP^* fate-mapping mice (**B**).

**Supplemental Figure S6.**
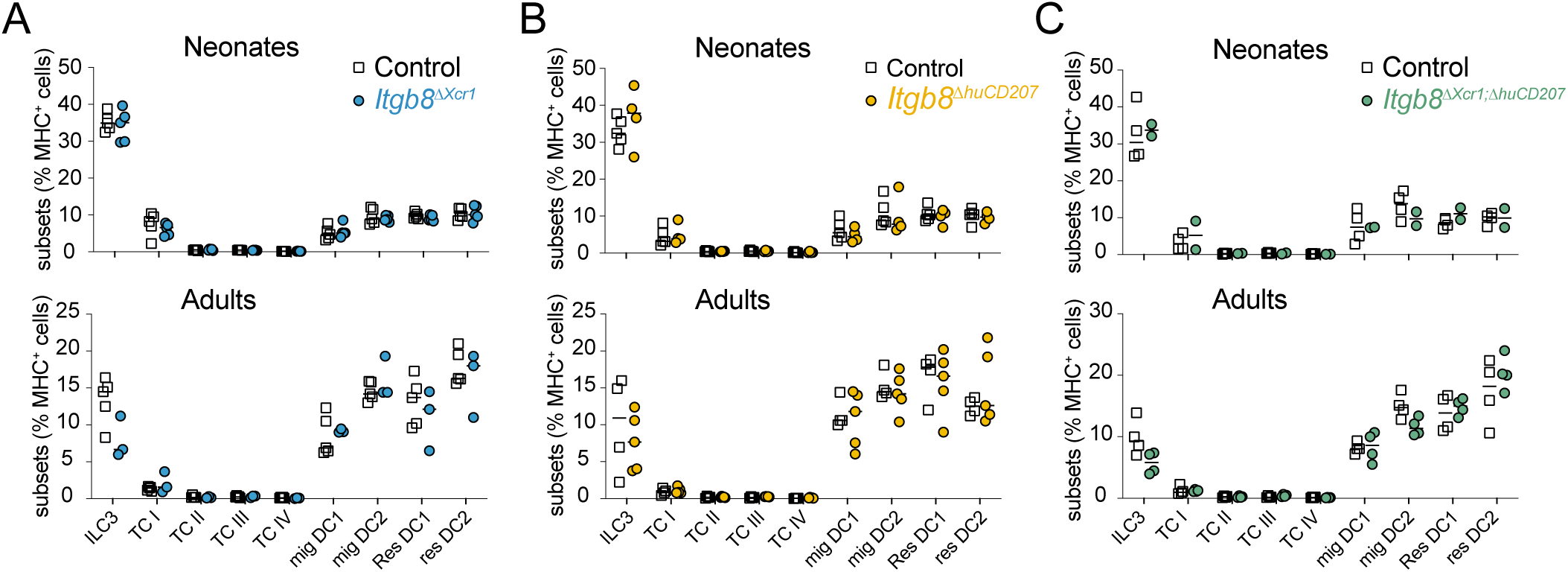
Frequency of MLN APCs in single (*Itgb8^ΔXcr^*^1^ and *Itgb8^ΔhuCD^*^207^) or double knockout (*Itgb8^ΔXcr^*^1^*^:huCD^*^207^) mice compared to littermate controls. **A-C.** Proportion of each APC subsets (gated as in Supplemental Figure S4) from the MLNs of 2-week-old neonatal (top) and adult (13 to 18-week-old, top) of *Itgb8^ΔXcr^*^1^(**A**), *Itgb8^ΔhuCD^*^207^ (**B**) and *Itgb8^ΔXcr^*^1^*^;ΔhuCD^*^207^ DKO mice (**C**). n=2 to 5 per group.

**Supplemental Figure S7.**
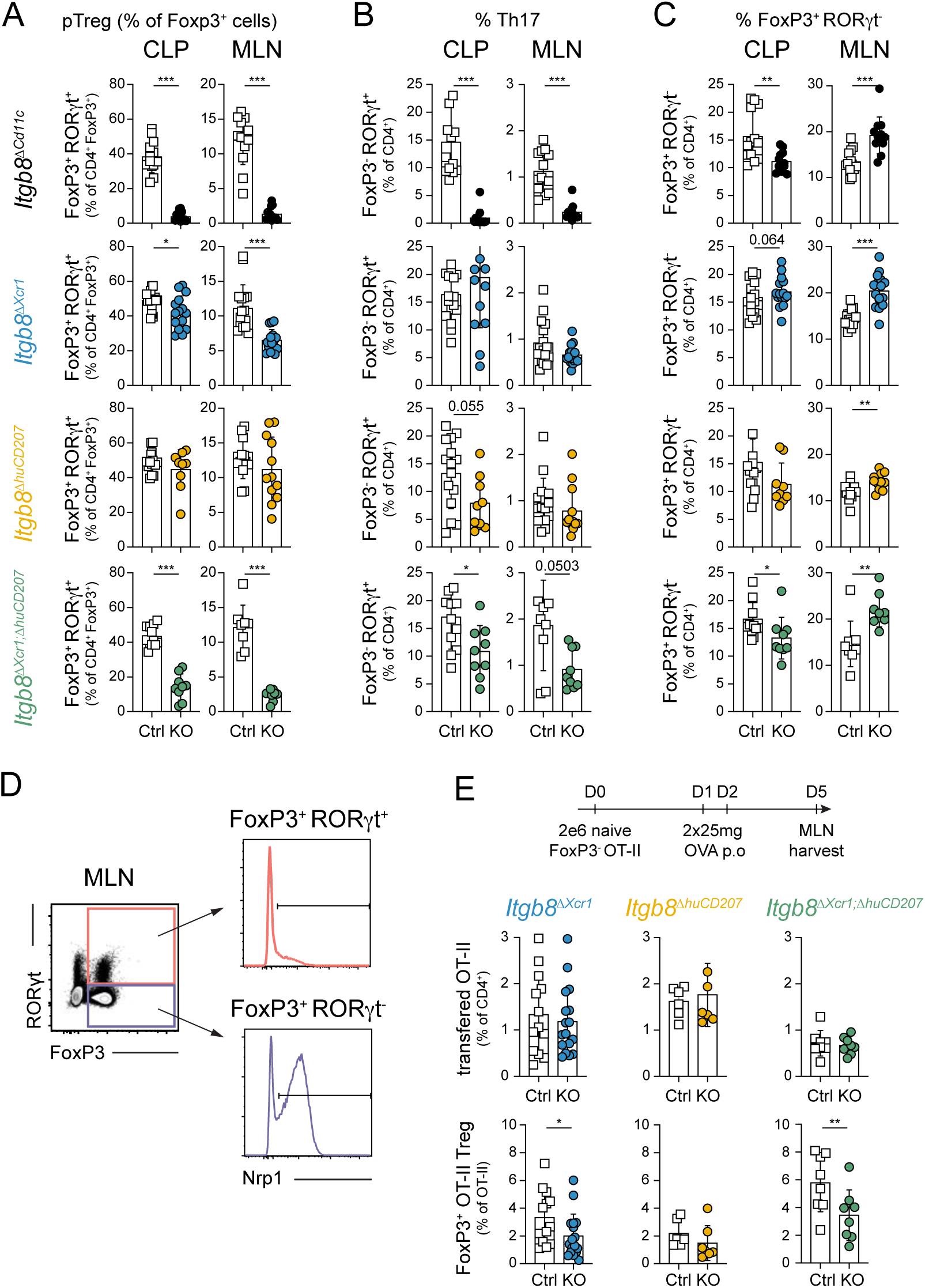
Frequency of Th17 and tTreg cells in β8-deficient mouse models. **A-C**. Summary graphs for frequencies of (**A**) RORγT^+^ pTreg, expressed as the% of all Foxp3^+^ CD4^+^ T cells and of Th17 (FoxP3^-^RORγt^+^) (**B**) and tTreg (FoxP3^+^RORγt^-^) (**C**) cells among total CD4^+^ T cells (**B-C**) in colonic lamina propria (CLP) and MLNs of *Itgb8^ΔCd^*^11c^ (black circles), *Itgb8^ΔXcr^*^1^ (blue circles), *Itgb8^ΔhuCD^*^207^ (yellow circles) and *Itgb8^ΔXcr^*^1^*^;ΔhuCD^*^207^ DKO (green circles), compared to littermate control mice (open squares). P values are reported as the following: *P < 0.05; **P < 0.01; ***P < 0.001; ****P < 0.0001. D. **E**. Experimental schematic (top) and quantification of the recovery of transferred OT-II T cells (middle) and of the *de novo* generation of Foxp3^+^ OT-II cell subsets (bottom) upon oral ovalbumine (OVA) administration: mice were adoptively transferred with FACS-sorted naïve Foxp3^-^OT-II cells on D0 before receiving two doses of i.g. OVA on D1 et D2. MLNs were harvested on D5. n=8. Error bars represent S.D. *P < 0.05; **P < 0.01; ***P < 0.001.

**Supplemental Figure S8.**
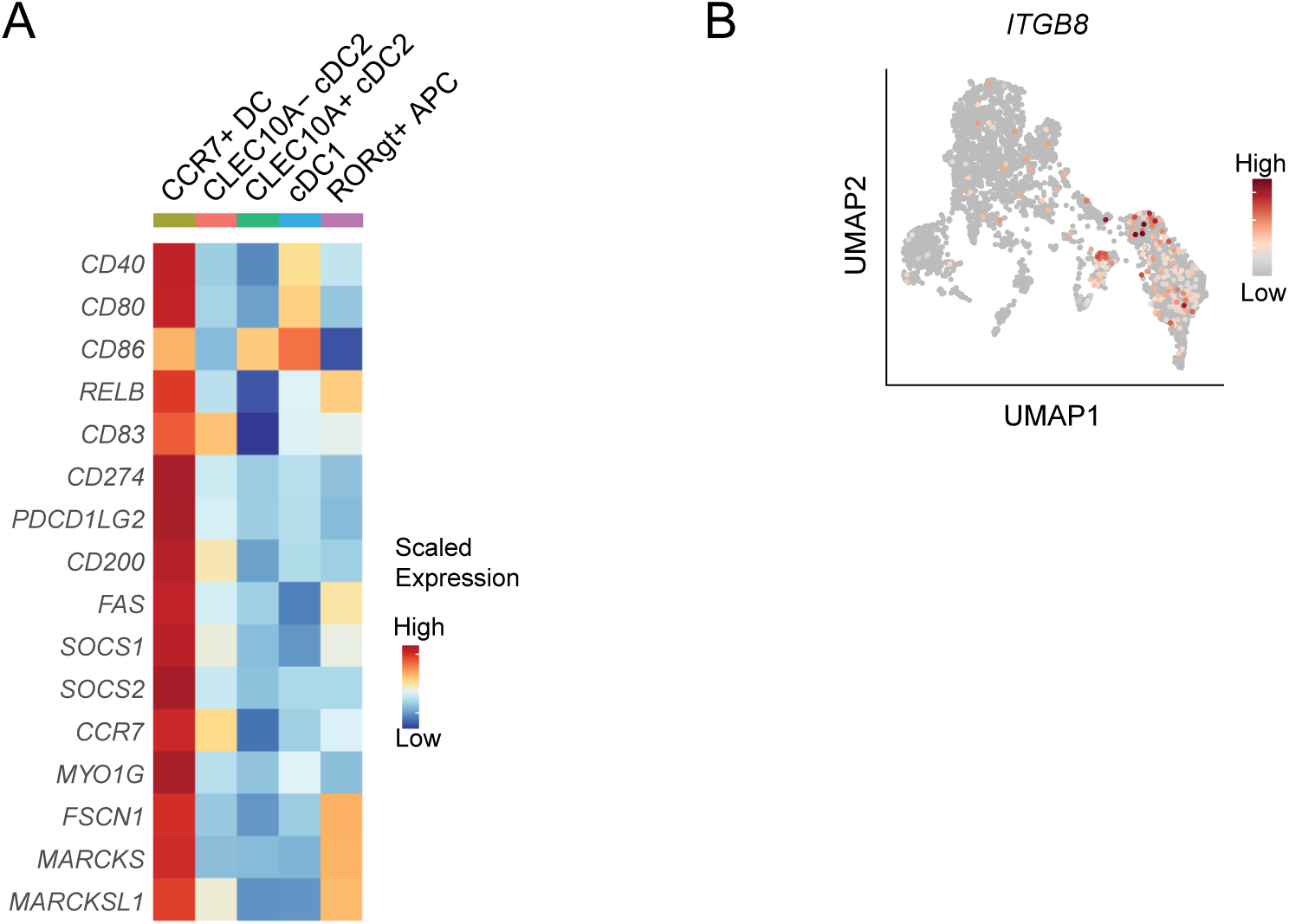
scRNAseq analysis of human MLN APCs. **A.** Heatmap showing expression of genes related to DC maturation and migration in human MLN APC subsets. **B**. UMAP of *ITGB8* expression in human MLN APCs. Cells are plotted in the order of expression intensity with highest expression in front.

**Suplementary Table 1.**
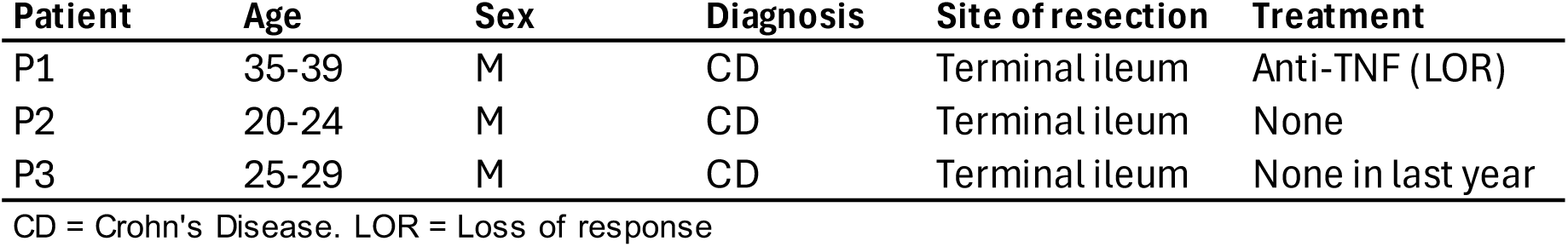
Patients’ characteristics.

## References

1. Maloy, K.J., and Powrie, F. (2011). Intestinal homeostasis and its breakdown in inflammatory bowel disease. Nature 474, 298–306. 10.1038/nature10208.

2. Caruso, R., Lo, B.C., and Núñez, G. (2020). Host–microbiota interactions in inflammatory bowel disease. Nat Rev Immunol 20, 411–426. 10.1038/s41577-019-0268-7.

3. Littman, D.R., and Rudensky, A.Y. (2010). Th17 and Regulatory T Cells in Mediating and Restraining Inflammation. Cell 140, 845–858. 10.1016/j.cell.2010.02.021.

4. Whibley, N., Tucci, A., and Powrie, F. (2019). Regulatory T cell adaptation in the intestine and skin. Nat Immunol 20, 386–396. 10.1038/s41590-019-0351-z.

5. Haribhai, D., Williams, J.B., Jia, S., Nickerson, D., Schmitt, E.G., Edwards, B., Ziegelbauer, J., Yassai, M., Li, S.-H., Relland, L.M., et al. (2011). A Requisite Role for Induced Regulatory T Cells in Tolerance Based on Expanding Antigen Receptor Diversity. Immunity 35, 109–122. 10.1016/j.immuni.2011.03.029.

6. Lathrop, S.K., Bloom, S.M., Rao, S.M., Nutsch, K., Lio, C.-W., Santacruz, N., Peterson, D.A., Stappenbeck, T.S., and Hsieh, C.-S. (2011). Peripheral education of the immune system by colonic commensal microbiota. Nature 478, 250–254. 10.1038/nature10434.

7. Izcue, A., Coombes, J.L., and Powrie, F. (2009). Regulatory Lymphocytes and Intestinal Inflammation. Annu. Rev. Immunol. 27, 313–338. 10.1146/annurev.immunol.021908.132657.

8. Xavier, R.J., and Podolsky, D.K. (2007). Unravelling the pathogenesis of inflammatory bowel disease. Nature 448, 427–434. 10.1038/nature06005.

9. Scott, C.L., Aumeunier, A.M., and Mowat, A.McI. (2011). Intestinal CD103+ dendritic cells: master regulators of tolerance? Trends in Immunology 32, 412–419. 10.1016/j.it.2011.06.003.

10. Coombes, J.L., Siddiqui, K.R.R., Arancibia-Cárcamo, C.V., Hall, J., Sun, C.-M., Belkaid, Y., and Powrie, F. (2007). A functionally specialized population of mucosal CD103+ DCs induces Foxp3+ regulatory T cells via a TGF-β– and retinoic acid–dependent mechanism. Journal of Experimental Medicine 204, 1757–1764. 10.1084/jem.20070590.

11. Sun, I.-H., Qualls, A.E., Yin, H.S., Wang, J., Arvedson, M.P., Germino, J., Horner, N.K., Zhong, S., Du, J., Valdearcos, M., et al. (2025). RORγt eTACs mediate oral tolerance and Treg induction. Journal of Experimental Medicine 222, e20250573. 10.1084/jem.20250573.

12. Worthington, J.J., Klementowicz, J.E., and Travis, M.A. (2011). TGFβ: a sleeping giant awoken by integrins. Trends in Biochemical Sciences 36, 47–54. 10.1016/j.tibs.2010.08.002.

13. Païdassi, H., Acharya, M., Zhang, A., Mukhopadhyay, S., Kwon, M., Chow, C., Stuart, L.M., Savill, J., and Lacy–Hulbert, A. (2011). Preferential Expression of Integrin αvβ8 Promotes Generation of Regulatory T Cells by Mouse CD103+ Dendritic Cells. Gastroenterology 141, 1813–1820. 10.1053/j.gastro.2011.06.076.

14. Worthington, J.J., Czajkowska, B.I., Melton, A.C., and Travis, M.A. (2011). Intestinal Dendritic Cells Specialize to Activate Transforming Growth Factor-β and Induce Foxp3+ Regulatory T Cells via Integrin αvβ8. Gastroenterology 141, 1802–1812. 10.1053/j.gastro.2011.06.057.

15. Worthington, J.J., Fenton, T.M., Czajkowska, B.I., Klementowicz, J.E., and Travis, M.A. (2012). Regulation of TGFβ in the immune system: An emerging role for integrins and dendritic cells. Immunobiology 217, 1259–1265. 10.1016/j.imbio.2012.06.009.

16. Travis, M.A., Reizis, B., Melton, A.C., Masteller, E., Tang, Q., Proctor, J.M., Wang, Y., Bernstein, X., Huang, X., Reichardt, L.F., et al. (2007). Loss of integrin αvβ8 on dendritic cells causes autoimmunity and colitis in mice. Nature 446, 361–365. 10.1038/nature06110.

17. Lacy-Hulbert, A., Smith, A.M., Tissire, H., Barry, M., Crowley, D., Bronson, R.T., Roes, J.T., Savill, J.S., and Hynes, R.O. (2007). Ulcerative colitis and autoimmunity induced by loss of myeloid αv integrins. Proceedings of the National Academy of Sciences 104, 15823–15828. 10.1073/pnas.0707421104.

18. Sun, T., Nguyen, A., and Gommerman, J.L. (2020). Dendritic Cell Subsets in Intestinal Immunity and Inflammation. The Journal of Immunology 204, 1075–1083. 10.4049/jimmunol.1900710.

19. Luda, K.M., Joeris, T., Persson, E.K., Rivollier, A., Demiri, M., Sitnik, K.M., Pool, L., Holm, J.B., Melo-Gonzalez, F., Richter, L., et al. (2016). IRF8 Transcription-Factor-Dependent Classical Dendritic Cells Are Essential for Intestinal T Cell Homeostasis. Immunity 44, 860–874. 10.1016/j.immuni.2016.02.008.

20. Hildner, K., Edelson, B.T., Purtha, W.E., Diamond, M., Matsushita, H., Kohyama, M., Calderon, B., Schraml, B.U., Unanue, E.R., Diamond, M.S., et al. (2008). Batf3 Deficiency Reveals a Critical Role for CD8α+ Dendritic Cells in Cytotoxic T Cell Immunity. Science 322, 1097–1100. 10.1126/science.1164206.

21. Brown, C.C., Gudjonson, H., Pritykin, Y., Deep, D., Lavallée, V.-P., Mendoza, A., Fromme, R., Mazutis, L., Ariyan, C., Leslie, C., et al. (2019). Transcriptional Basis of Mouse and Human Dendritic Cell Heterogeneity. Cell 176, 846–863.e24. 10.1016/j.cell.2019.09.035.

22. Caton, M.L., Smith-Raska, M.R., and Reizis, B. (2007). Notch–RBP-J signaling controls the homeostasis of CD8− dendritic cells in the spleen. Journal of Experimental Medicine 204, 1653–1664. 10.1084/jem.20062648.

23. Satpathy, A.T., Briseño, C.G., Lee, J.S., Ng, D., Manieri, N.A., Kc, W., Wu, X., Thomas, S.R., Lee, W.-L., Turkoz, M., et al. (2013). Notch2-dependent classical dendritic cells orchestrate intestinal immunity to attaching-and-effacing bacterial pathogens. Nat Immunol 14, 937–948. 10.1038/ni.2679.

24. Lewis, K.L., Caton, M.L., Bogunovic, M., Greter, M., Grajkowska, L.T., Ng, D., Klinakis, A., Charo, I.F., Jung, S., Gommerman, J.L., et al. (2011). Notch2 Receptor Signaling Controls Functional Differentiation of Dendritic Cells in the Spleen and Intestine. Immunity 35, 780–791. 10.1016/j.immuni.2011.08.013.

25. Persson, E.K., Uronen-Hansson, H., Semmrich, M., Rivollier, A., Hägerbrand, K., Marsal, J., Gudjonsson, S., Håkansson, U., Reizis, B., Kotarsky, K., et al. (2013). IRF4 Transcription-Factor-Dependent CD103+CD11b+ Dendritic Cells Drive Mucosal T Helper 17 Cell Differentiation. Immunity 38, 958–969. 10.1016/j.immuni.2013.03.009.

26. Welty, N.E., Staley, C., Ghilardi, N., Sadowsky, M.J., Igyártó, B.Z., and Kaplan, D.H. (2013). Intestinal lamina propria dendritic cells maintain T cell homeostasis but do not affect commensalism. Journal of Experimental Medicine 210, 2011–2024. 10.1084/jem.20130728.

27. Esterházy, D., Loschko, J., London, M., Jove, V., Oliveira, T.Y., and Mucida, D. (2016). Classical dendritic cells are required for dietary antigen–mediated induction of peripheral Treg cells and tolerance. Nat Immunol 17, 545–555. 10.1038/ni.3408.

28. Boucard-Jourdin, M., Kugler, D., Endale Ahanda, M.-L., This, S., De Calisto, J., Zhang, A., Mora, J.R., Stuart, L.M., Savill, J., Lacy-Hulbert, A., et al. (2016). β8 Integrin Expression and Activation of TGF-β by Intestinal Dendritic Cells Are Determined by Both Tissue Microenvironment and Cell Lineage. J Immunol 167, 1968–1978. 10.4049/jimmunol.1600244.

29. Akagbosu, B., Tayyebi, Z., Shibu, G., Paucar Iza, Y.A., Deep, D., Parisotto, Y.F., Fisher, L., Pasolli, H.A., Thevin, V., Elmentaite, R., et al. (2022). Novel antigen-presenting cell imparts Treg-dependent tolerance to gut microbiota. Nature 610, 752–760. 10.1038/s41586-022-05309-5.

30. Kedmi, R., Najar, T.A., Mesa, K.R., Grayson, A., Kroehling, L., Hao, Y., Hao, S., Pokrovskii, M., Xu, M., Talbot, J., et al. (2022). A RORγt+ cell instructs gut microbiota-specific Treg cell differentiation. Nature 610, 737–743. 10.1038/s41586-022-05089-y.

31. Lyu, M., Suzuki, H., Kang, L., Gaspal, F., Zhou, W., Goc, J., Zhou, L., Zhou, J., Zhang, W., Shen, Z., et al. (2022). ILC3s select microbiota-specific regulatory T cells to establish tolerance in the gut. Nature 610, 744–751. 10.1038/s41586-022-05141-x.

32. Schraml, B.U., van Blijswijk, J., Zelenay, S., Whitney, P.G., Filby, A., Acton, S.E., Rogers, N.C., Moncaut, N., Carvajal, J.J., and Reis e Sousa, C. (2013). Genetic Tracing via DNGR-1 Expression History Defines Dendritic Cells as a Hematopoietic Lineage. Cell 154, 843–858. 10.1016/j.cell.2013.07.014.

33. Salvermoser, J., van Blijswijk, J., Papaioannou, N.E., Rambichler, S., Pasztoi, M., Pakalniškytė, D., Rogers, N.C., Keppler, S.J., Straub, T., Reis e Sousa, C., et al. (2018). Clec9a-Mediated Ablation of Conventional Dendritic Cells Suggests a Lymphoid Path to Generating Dendritic Cells In Vivo. Front. Immunol. 9. 10.3389/fimmu.2018.00699.

34. Nakawesi, J., This, S., Hütter, J., Boucard-Jourdin, M., Barateau, V., Muleta, K.G., Gooday, L.J., Thomsen, K.F., López, A.G., Ulmert, I., et al. (2021). αvβ8 integrin-expression by BATF3-dependent dendritic cells facilitates early IgA responses to Rotavirus. Mucosal Immunology 14, 53–67. 10.1038/s41385-020-0276-8.

35. Wang, J., Lareau, C.A., Bautista, J.L., Gupta, A.R., Sandor, K., Germino, J., Yin, Y., Arvedson, M.P., Reeder, G.C., Cramer, N.T., et al. (2021). Single-cell multiomics defines tolerogenic extrathymic Aire-expressing populations with unique homology to thymic epithelium. Science Immunology 6, eabl5053. 10.1126/sciimmunol.abl5053.

36. Park, T., Leslie, C., Rudensky, A.Y., and Brown, C.C. (2023). Reconciling the spectrum of RORγt+ antigen-presenting cells. Preprint at bioRxiv, 10.1101/2023.11.01.565227 https://doi.org/10.1101/2023.11.01.565227.

37. Narasimhan, H., Richter, M.L., Shakiba, R., Papaioannou, N.E., Stehle, C., Ravi Rengarajan, K., Ulmert, I., Kendirli, A., de la Rosa, C., Kuo, P.-Y., et al. (2025). RORγt-expressing dendritic cells are functionally versatile and evolutionarily conserved antigen-presenting cells. Proceedings of the National Academy of Sciences 122, e2417308122. 10.1073/pnas.2417308122.

38. Fu, L., Upadhyay, R., Pokrovskii, M., Chen, F.M., Romero-Meza, G., Griesemer, A., and Littman, D.R. (2025). PRDM16-dependent antigen-presenting cells induce tolerance to gut antigens. Nature, 1–10. 10.1038/s41586-025-08982-4.

39. Rodrigues, P.F., Wu, S., Trsan, T., Panda, S.K., Fachi, J.L., Liu, Y., Du, S., Oliveira, S. de, Antonova, A.U., Khantakova, D., et al. (2025). Rorγt-positive dendritic cells are required for the induction of peripheral regulatory T cells in response to oral antigens. Cell 188, 2720–2737.e22. 10.1016/j.cell.2025.03.020.

40. Abram, C.L., Roberge, G.L., Hu, Y., and Lowell, C.A. (2014). Comparative analysis of the efficiency and specificity of myeloid-Cre deleting strains using ROSA-EYFP reporter mice. Journal of Immunological Methods 408, 89–100. 10.1016/j.jim.2014.05.009.

41. Mattiuz, R., Wohn, C., Ghilas, S., Ambrosini, M., Alexandre, Y.O., Sanchez, C., Fries, A., Vu Manh, T.-P., Malissen, B., Dalod, M., et al. (2018). Novel Cre-Expressing Mouse Strains Permitting to Selectively Track and Edit Type 1 Conventional Dendritic Cells Facilitate Disentangling Their Complexity in vivo. Front. Immunol. 9. 10.3389/fimmu.2018.02805.

42. Kaplan, D.H., Li, M.O., Jenison, M.C., Shlomchik, W.D., Flavell, R.A., and Shlomchik, M.J. (2007). Autocrine/paracrine TGFβ1 is required for the development of epidermal Langerhans cells. Journal of Experimental Medicine 204, 2545–2552. 10.1084/jem.20071401.

43. Weiss, J.M., Bilate, A.M., Gobert, M., Ding, Y., Curotto de Lafaille, M.A., Parkhurst, C.N., Xiong, H., Dolpady, J., Frey, A.B., Ruocco, M.G., et al. (2012). Neuropilin 1 is expressed on thymus-derived natural regulatory T cells, but not mucosa-generated induced Foxp3+ T reg cells. J Exp Med 204, 1723–1742. 10.1084/jem.20120914.

44. Yadav, M., Louvet, C., Davini, D., Gardner, J.M., Martinez-Llordella, M., Bailey-Bucktrout, S., Anthony, B.A., Sverdrup, F.M., Head, R., Kuster, D.J., et al. (2012). Neuropilin-1 distinguishes natural and inducible regulatory T cells among regulatory T cell subsets in vivo. J Exp Med 209, 1713–1722. 10.1084/jem.20120822.

45. Melton, A.C., Bailey-Bucktrout, S.L., Travis, M.A., Fife, B.T., Bluestone, J.A., and Sheppard, D. (2010). Expression of αvβ8 integrin on dendritic cells regulates Th17 cell development and experimental autoimmune encephalomyelitis in mice. J Clin Invest 120, 4436–4444. 10.1172/JCI43786.

46. Acharya, M., Mukhopadhyay, S., Païdassi, H., Jamil, T., Chow, C., Kissler, S., Stuart, L.M., Hynes, R.O., and Lacy-Hulbert, A. (2010). αv Integrin expression by DCs is required for Th17 cell differentiation and development of experimental autoimmune encephalomyelitis in mice. J Clin Invest 120, 4445–4452. 10.1172/JCI43796.

47. Campos Canesso, M.C., de Castro, T.B.R., Nakandakari-Higa, S., Lockhart, A., Luehr, J., Bortolatto, J., Parsa, R., Esterházy, D., Lyu, M., Liu, T.-T., et al. (2024). Identification of antigen-presenting cell–T cell interactions driving immune responses to food. Science 387, eado5088. 10.1126/science.ado5088.

48. Rudnitsky, A., Oh, H., Margolin, M., Dassa, B., Shteinberg, I., Stoler-Barak, L., Shulman, Z., and Kedmi, R. (2025). A coordinated cellular network regulates tolerance to food. Nature 644, 231–240. 10.1038/s41586-025-09173-x.

49. Cabric, V., Franco Parisotto, Y., Park, T., Akagbosu, B., Zhao, Z., Lo, Y., Shibu, G., Fisher, L., Paucar Iza, Y.A., Leslie, C., et al. (2025). A wave of Thetis cells imparts tolerance to food antigens early in life. Science 0, eadp0535. 10.1126/science.adp0535.

50. Fenton, T.M., Kelly, A., Shuttleworth, E.E., Smedley, C., Atakilit, A., Powrie, F., Campbell, S., Nishimura, S.L., Sheppard, D., Levison, S., et al. (2017). Inflammatory cues enhance TGFβ activation by distinct subsets of human intestinal dendritic cells via integrin αvβ8. Mucosal Immunology 10, 624–634. 10.1038/mi.2016.94.

51. Nutsch, K., Chai, J.N., Ai, T.L., Russler-Germain, E., Feehley, T., Nagler, C.R., and Hsieh, C.-S. (2016). Rapid and Efficient Generation of Regulatory T Cells to Commensal Antigens in the Periphery. Cell Reports 17, 206–220. 10.1016/j.celrep.2016.08.092.

52. Mucida, D., Park, Y., Kim, G., Turovskaya, O., Scott, I., Kronenberg, M., and Cheroutre, H. (2007). Reciprocal TH17 and Regulatory T Cell Differentiation Mediated by Retinoic Acid. Science 317, 256–260. 10.1126/science.1145697.

53. Idoyaga, J., Fiorese, C., Zbytnuik, L., Lubkin, A., Miller, J., Malissen, B., Mucida, D., Merad, M., and Steinman, R.M. (2013). Specialized role of migratory dendritic cells in peripheral tolerance induction. J Clin Invest 123, 844–854. 10.1172/JCI65260.

54. Dalod, M., and Scheu, S. (2022). Dendritic cell functions in vivo: A user’s guide to current and next-generation mutant mouse models. European Journal of Immunology 52, 1712–1749. 10.1002/eji.202149513.

55. Lança, T., Ungerbäck, J., Silva, C.D., Joeris, T., Ahmadi, F., Vandamme, J., Svensson-Frej, M., Mowat, A.M., Kotarsky, K., Sigvardsson, M., et al. (2022). IRF8 deficiency induces the transcriptional, functional, and epigenetic reprogramming of cDC1 into the cDC2 lineage. Immunity 55, 1431–1447.e11. 10.1016/j.immuni.2022.06.006.

56. Tussiwand, R., Lee, W.-L., Murphy, T.L., Mashayekhi, M., Kc, W., Albring, J.C., Satpathy, A.T., Rotondo, J.A., Edelson, B.T., Kretzer, N.M., et al. (2012). Compensatory dendritic cell development mediated by BATF–IRF interactions. Nature 490, 502–507. 10.1038/nature11531.

57. Fenton, T.M., Wulff, L., Väänänen, V., Jones, G.-R., Lemvigh, C.K., Riis, L.B., Wewer, M.D., Vandamme, J., Jørgensen, P.B., Bain, C.C., et al. (2025). Heterogeneity of the intestinal mononuclear phagocyte compartment in health and inflammatory bowel disease. Science Immunology 10, eadz8650. 10.1126/sciimmunol.adz8650.

58. de Lange, K.M., Moutsianas, L., Lee, J.C., Lamb, C.A., Luo, Y., Kennedy, N.A., Jostins, L., Rice, D.L., Gutierrez-Achury, J., Ji, S.-G., et al. (2017). Genome-wide association study implicates immune activation of multiple integrin genes in inflammatory bowel disease. Nat Genet 49, 256–261. 10.1038/ng.3760.

59. Okada, Y., Tsuzuki, Y., Takeshi, T., Furuhashi, H., Higashiyama, M., Watanabe, C., Shirakabe, K., Kurihara, C., Komoto, S., Tomita, K., et al. (2018). Novel probiotics isolated from a Japanese traditional fermented food, Funazushi, attenuates DSS-induced colitis by increasing the induction of high integrin αv/β8-expressing dendritic cells. J Gastroenterol 53, 407–418. 10.1007/s00535-017-1362-x.

60. Minagawa, S., Lou, J., Seed, R.I., Cormier, A., Wu, S., Cheng, Y., Murray, L., Tsui, P., Connor, J., Herbst, R., et al. (2014). Selective Targeting of TGF-β Activation to Treat Fibroinflammatory Airway Disease. Science Translational Medicine 6, 241ra79–241ra79. 10.1126/scitranslmed.3008074.

61. Ulezko Antonova, A., Lonardi, S., Monti, M., Missale, F., Fan, C., Coates, M.L., Bugatti, M., Jaeger, N., Fernandes Rodrigues, P., Brioschi, S., et al. (2023). A distinct human cell type expressing MHCII and RORγt with dual characteristics of dendritic cells and type 3 innate lymphoid cells. Proceedings of the National Academy of Sciences 120, e2318710120. 10.1073/pnas.2318710120.

62. Proctor, J.M., Zang, K., Wang, D., Wang, R., and Reichardt, L.F. (2005). Vascular Development of the Brain Requires β8 Integrin Expression in the Neuroepithelium. J. Neurosci. 25, 9940–9948. 10.1523/JNEUROSCI.3467-05.2005.

63. Srinivas, S., Watanabe, T., Lin, C.-S., William, C.M., Tanabe, Y., Jessell, T.M., and Costantini, F. (2001). Cre reporter strains produced by targeted insertion of EYFP and ECFP into the ROSA26 locus. BMC Developmental Biology 1, 4. 10.1186/1471-213X-1-4.

64. Barnden, M.J., Allison, J., Heath, W.R., and Carbone, F.R. (1998). Defective TCR expression in transgenic mice constructed using cDNA-based α- and β-chain genes under the control of heterologous regulatory elements. Immunology & Cell Biology 76, 34.–40. 10.1046/j.1440-1711.1998.00709.x.

65. Wang, Y., Kissenpfennig, A., Mingueneau, M., Richelme, S., Perrin, P., Chevrier, S., Genton, C., Lucas, B., DiSanto, J.P., Acha-Orbea, H., et al. (2008). Th2 Lymphoproliferative Disorder of LatY136F Mutant Mice Unfolds Independently of TCR-MHC Engagement and Is Insensitive to the Action of Foxp3+ Regulatory T Cells1. The Journal of Immunology 180, 1565–1575. 10.4049/jimmunol.180.3.1565.

66. Koelink, P.J., Wildenberg, M.E., Stitt, L.W., Feagan, B.G., Koldijk, M., van ‘t Wout, A.B., Atreya, R., Vieth, M., Brandse, J.F., Duijst, S., et al. (2018). Development of Reliable, Valid and Responsive Scoring Systems for Endoscopy and Histology in Animal Models for Inflammatory Bowel Disease. Journal of Crohn’s and Colitis 12, 794–803. 10.1093/ecco-jcc/jjy035.

67. Hao, Y., Stuart, T., Kowalski, M.H., Choudhary, S., Hoffman, P., Hartman, A., Srivastava, A., Molla, G., Madad, S., Fernandez-Granda, C., et al. (2024). Dictionary learning for integrative, multimodal and scalable single-cell analysis. Nat Biotechnol 42, 293–304. 10.1038/s41587-023-01767-y.

68. Luecken, M.D., and Theis, F.J. (2019). Current best practices in single-cell RNA-seq analysis: a tutorial. Mol Syst Biol 15, MSB188746. 10.15252/msb.20188746.

69. Germain, P.-L., Lun, A., Meixide, C.G., Macnair, W., and Robinson, M.D. (2022). Doublet identification in single-cell sequencing data using *scDblFinder*. Preprint at F1000Research, 10.12688/f1000research.73600.2 https://doi.org/10.12688/f1000research.73600.2.

70. Korsunsky, I., Millard, N., Fan, J., Slowikowski, K., Zhang, F., Wei, K., Baglaenko, Y., Brenner, M., Loh, P., and Raychaudhuri, S. (2019). Fast, sensitive and accurate integration of single-cell data with Harmony. Nat Methods 16, 1289–1296. 10.1038/s41592-019-0619-0.

